# Two cortical mechanisms for natural audiovisual processing

**DOI:** 10.1101/2025.11.05.686819

**Authors:** Subha Nawer Pushpita, Leila Wehbe

## Abstract

Understanding how the human brain processes natural audiovisual information remains a central challenge in cognitive neuroscience. Progress has been limited by the difficulty of modeling complex audiovisual stimuli; prior work has therefore relied largely on short, controlled, or unimodal stimuli, leaving cortical mechanisms supporting real-world comprehension poorly characterized. Although recent AI advances enable extraction of high-dimensional, time-resolved features from naturalistic stimuli, how cortical regions dynamically prioritize auditory and visual information as time unfolds remains largely unexplored. Using large-scale fMRI data collected during movie viewing, we developed two complementary prediction-based approaches to map moment-by-moment sensory processing across cortex: one detects sustained periods when one modality predicts a region better than the other, identifying regions that switch the modality they encode for meaningful stretches of time; while the other identifies periods when both modalities predict the region well, revealing balanced audiovisual representations. Together, these analyses reveal two cortical patterns of audiovisual processing: two modality-switching “bows” (a posterior bow encircling category-selective visual cortex and an anterior bow spanning dorsolateral frontal cortex) and an arrow-like axis jointly representing both modalities (extending from lateral occipital into temporal cortex). We further show that switching is not driven by reduced low-level information in the non-dominant modality, but by reduced cortical encoding of that information, suggesting active sensory reweighting. Modality dominance also cannot be explained simply by the dominant modality conveying a region’s preferred concepts; switching regions do not merely track whichever modality expresses these concepts. Together, these systems suggest a cortical architecture that adaptively reweights sensory inputs while maintaining balanced multimodal representations during naturalistic comprehension.

## Main

Humans effortlessly navigate a rich multimodal world, comprehend complex continuous stream of audiovisual stimuli like movies and TV series, and seamlessly integrate rich information conveyed across multiple sensory modalities. However, the vast majority of cognitive neuroscience experiments, even those using naturalistic stimuli, focus on one modality at a time, such as vision [1, 2, 3, 4] or audition/language processing [5, 6, 7, 8], not studying how these modalities are used together in everyday life. There are a few prevailing theories of how the brain supports multimodal comprehension. The hub-and-spoke view holds that modality-specific “spokes” in sensory cortices converge on an amodal hub in the anterior temporal lobe (ATL), consistent with findings that ATL degeneration yields semantic dementia while syntactical, numerical abilities, and executive functions are fairly intact [9, 10, 11, 12]. In contrast, the convergence-zone view proposes that modality-specific and amodal codes integrate across multiple cortical sites beyond ATL[13]. Extending beyond these models, Driver and Noesselt [14] emphasized that multisensory integration itself can occur at multiple hierarchical levels - from direct interactions between unisensory cortices, to specialized multisensory patches, to higher-order hubs that flexibly coordinate and reshape the balance between unisensory and multisensory representations. Crucially, border regions such as angular gyrus, precuneus, and middle temporal gyrus encode the same semantic categories across visual and linguistic input [15, 16]. There has also been evidence for the semantic alignment hypothesis, suggesting that each location on the anterior edge of the visual cortex that encodes a particular visual category is followed directly anteriorly by a region that encodes the same category in linguistic form [17]. In parallel, distinct patches in superior temporal and parietal cortex were shown to represent social interactions—an intrinsically multimodal construct—highlighting that the brain’s capacity for multimodal comprehension extends beyond objects and semantics to the dynamic understanding of how people relate to one another [18]. Beauchamp’s work demonstrated that the superior temporal sulcus integrates auditory and visual inputs, with responses exceeding those to either modality alone, establishing it as a key cortical locus for the comprehension of audiovisual speech and action [19].

However, prior work has primarily examined the anatomical loci of multisensory integration and comprehension using primarily short, carefully controlled audiovisual stimuli, and often using non-simultaneous auditory and visual stimuli [19, 1, 2, 3, 4]. Far less is known about the neural mechanisms that govern how content demands at different moments within long, naturalistic audiovisual streams drive cortical regions to adopt diverse modes of audiovisual processing. Further, studies that do use naturalistic stimuli and analysis methods such as encoding models [20] or representation similarity analysis [21], focus on identifying fixed feature sensitivity of a given region, and do not evaluate whether the sensitivity of these regions changes in time - leading to implications that a region’s modality preference or integrative role is static. The assumption of representations being static is problematic for natural narratives, where the informational burden can shift dramatically between modalities over time.

To address this gap, we introduce a new hypothesis space (Figure 1) that incorporates the temporal dynamics of sensory processing: some regions show sustained unisensory dominance - favoring either visual (Figure 1a) or auditory (Figure 1b) inputs throughout (here, auditory refers broadly to all perceptual information received through the ears, including speech, language, prosody, and non-speech environmental sounds) - whereas others act as jointly predicted regions (Figure 1c), reflecting information from both modalities, potentially supporting integrative processing or amodal concept representation, and others emerge as “switching regions” that flexibly shift their preference (Figure 1d), significantly favoring auditory input at some moments and visual input at others.

**Figure 1:**
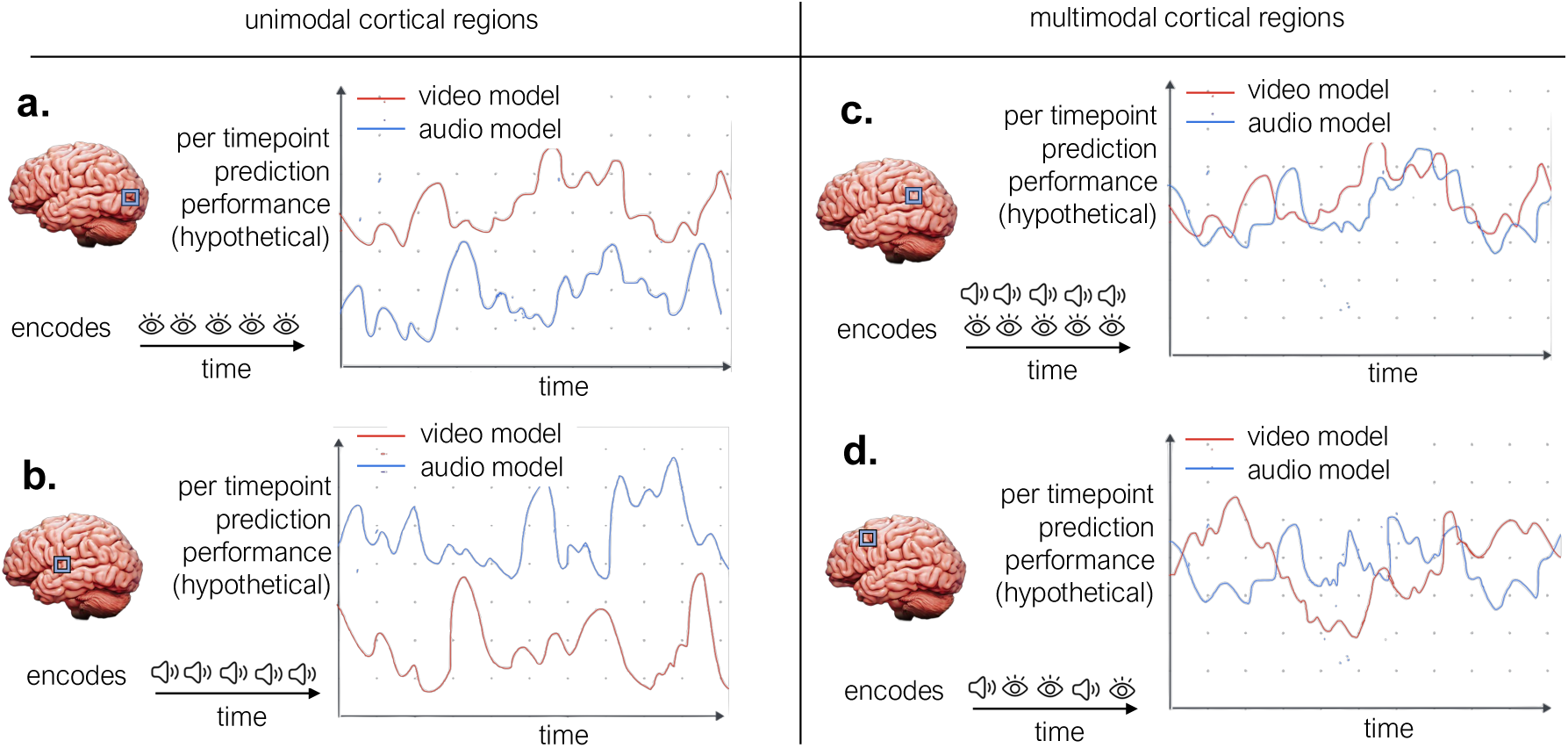
Hypothesis space for unimodal and multimodal regions. We illustrate four canonical patterns of audiovisual encoding that can explain regional differences in prediction performance. **(a)** In unimodal visual regions, BOLD responses will be predicted substantially better by models fitted on visual features (red) than by those fitted on auditory features (blue), reflecting stable visual selectivity. **(b)** Conversely, unimodal auditory/language regions are consistently better predicted by auditory feature–based models, reflecting stable auditory/language selectivity. **(c)** In some multimodal regions, BOLD responses are predicted well by both feature types throughout time, indicating concurrent encoding of auditory and visual information. **(d)** Other multimodal regions alternate between periods dominated by auditory versus visual feature prediction, suggesting dynamic switching of modality preference. Together, these four patterns capture the range of possible temporal dynamics by which cortical regions may support comprehension of naturalistic audiovisual stimuli.

To empirically test this hypothesis space, we adopted a data-driven approach using the Courtois NeuroMod dataset[22, 23], released as part of Algonauts 2025 challenge [23], which consists of fMRI BOLD responses from four human subjects to more than 36 hours of TV series and movies. The data for each participant has been averaged into 1000 cortical parcels from the Schaefer atlas [24], and we further average the cross-participant activity for each parcel. We fit one encoding model for each modality that predicts each parcel activity as a function of distinct auditory or visual feature sets (Figure 2a, and details in Methods). We extracted auditory features from an audio LLM, Qwen2 Audio-7B instruct [25]. This model is trained on large audio–text corpora for speech recognition, emotion, speaker and event identification, and higher-level instruction-following tasks such as transcription, translation, and spoken question answering. Therefore, the collected features capture a hierarchy of paralinguistic and semantic information-from low-level acoustic structure to higher-order features such as sentence meaning, speaker identity, affect, and communicative intent-providing a rich representation of the auditory and semantic content beyond the speech signal itself. We extracted visual features from Qwen 2.5 VL[26] and VideoMAE[27]. Qwen 2.5 VL is trained on large-scale image–text and video–text datasets to optimize object- and region-level grounding, temporal reasoning, and action recognition across short and long timescales, and multimodal instruction-following tasks including captioning, question answering, and event localization whereas VideoMAE is trained for fine-grained short term temporal understanding. Hence, the collected features encode a broad spectrum of visual information-from static object and scene semantics to dynamic motions and event structure-capturing both momentary visual details and temporally extended context relevant for interpreting naturalistic audiovisual stimuli. Note that the audio features are derived solely from the audio stream, whereas the visual features are computed solely from the video frames. Together, the audio and visual features can predict a large swathe of cortex with robust prediction performance (see Figure 2d and Extended Figure E2 for the components of the explained variance).

**Figure 2:**
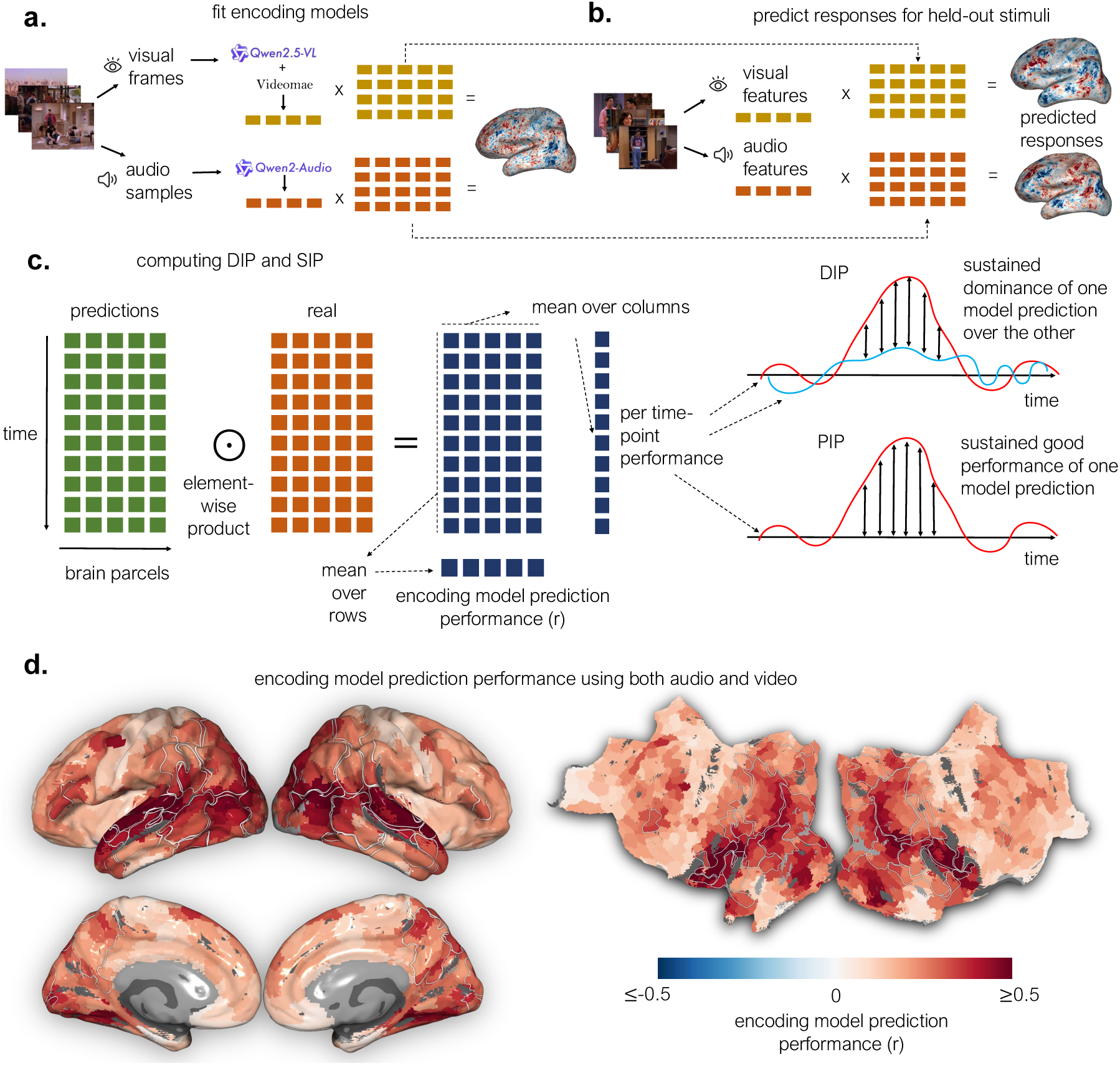
Overview of encoding model framework and computational paradigms. (a) We fit separate encoding models using features derived from the auditory stream (Qwen2-Audio) and the visual stream (VideoMAE and Qwen2.5-VL). The auditory model captures both acoustic and paralinguistic/languagerelated structure rather than only low-level sound features, while the visual model encodes spatiotemporal information and actions from video frames. Ridge regression was used to learn parcel-wise linear mappings between each feature space and measured BOLD responses (averaged across participants). (b) The trained models were then applied to predict BOLD activity for held-out movie segments, yielding time-resolved predictions from the visual and auditory models that could be compared across cortical regions. (c) We introduce two complementary data-driven approaches - the Dominance Indication Paradigm (DIP) and Performance Indication Paradigm (PIP), to characterize temporal dynamics of modality-specific and shared encoding. Instead of computing model performance across all timepoints for every parcel/voxel as in traditional encoding analyses, we quantified prediction performance at each timepoint within each region by taking the element-wise product between model predictions and observed responses, then averaging across parcels in that region. DIP captures periods of sustained dominance by one modality’s model over the other, whereas PIP captures periods of jointly strong performance by both models. (d) Parcel-wise prediction accuracy (Pearson correlation, *r*) of a concatenated audiovisual model on held-out stimuli. Both inflated (left) and flattened (left) cortical surfaces reveal strong prediction performance along posterior visual areas, extending anteriorly into lateral, ventral, dorsal and medial regions, spanning the temporal cortex and parts of the frontal cortex (see region names in Extended Figure E1). These widespread high-performing zones reflect the robust encoding of visual and auditory–linguistic features across large portions of cortex.

We introduce two complementary frameworks to track the temporal dynamics of modality engagement across the cortex: the Dominance Indication Paradigm (DIP) and the Performance Indication Paradigm (PIP). DIP marks timepoints where prediction accuracy on held-out data is substantially better accounted for by the audio-based model than by the video-based model (or vice versa), evaluated independently for each region, thereby identifying regions that alternate their dominant sensory modality over time (Figure 2c). PIP, in contrast, identifies time windows where either model robustly predicts the neural response—and particularly where both do so concurrently—computed separately for each region to reveal those regions that encode information from both modalities in parallel (Figure 2c). We further developed a specifically modified transformer model that predicts whole-brain fMRI responses while dynamically weighting audio and visual inputs for each cortical region, providing an automated means to analyze cortical regions by their modality-specific tuning profiles over time.

## Results

### Two Circular Bands of Regions Around Category-Selective Visual Cortex and Lateral Frontal Cortex Show Remarkable Switching

To operationalize DIP, we evaluated separately trained audio- and video-based encoding models on held-out movie data, quantifying differences in their prediction accuracy at each region of interest (ROI, defined using the Harvard–Oxford cortical atlas [28]) and time point (Figure 2a-b). Reliable differences between the time point performance of models should be large enough and sustained for a long enough period of time. Instead of picking these parameters arbitrarily, we adapted the threshold-free cluster enhancement method (TFCE) [29], applying it to performance differences standardized against the permutation null at each time point so that all time points contribute on a common scale. We identified TRs (fMRI time unit) for which the observed TFCE statistic exceeded, in at least 80% of block-wise permutations, the maximum TFCE statistic across all time points within that permutation. Comparing against the permutation-wise maximum rather than the time point-specific null controls the family-wise error rate across time points (details in Methods). Note that our characterization of switching regions does not qualitatively change when we change the threshold from 80%, as shown in Supplementary Figure S5. By conducting this analysis in both directions - testing when the audio-based model outperformed the video-based model and vice versa - we isolated timepoints where one modality’s model had a reliable advantage in predictive performance over the other, rather than merely a marginal difference in fit. This yielded, for each ROI, temporal segments of audio- or video-predictive dominance, providing a fine-grained measure of when and where cortical responses were better accounted for by one modality over the other.

Aggregating, for each region, the total duration of audio-dominant segments (*D*_audio_) and the total duration of video-dominant segments (*D*_video_) across all sessions revealed a striking organization of “switching” regions forming two distinct bow-like bands (Figure 3a): a posterior bow encircling visual cortex—including the medial precuneus, superior parietal lobule, posterior angular gyrus, anterior lateral occipital cortex, and temporo-occipital divisions of the middle and inferior temporal gyri—and a smaller anterior bow spanning dorso-lateral and dorso-medial frontal association areas, including parts of the superior, middle, and inferior frontal gyri. To determine which regions significantly alternated between modalities, we defined our switching index as *S* = minimum(*D*_audio_*, D*_video_), which allows us to identify regions that both show switching between the two modalities and switch for a large number of time points. This metric is valid for measuring switching in our setup because the distribution of dominant segment length in each switching region in Extended Figure (E5) indicates that dominant segments are intermediate in length, with the lengths mostly ranging from 10 to 18 TRs, or around 15 seconds to 27 seconds. Thus, each of the switching regions does display preference for each modality in multiple scenes, in contrast with spending a long time in one modality and then in the other. We conducted exhaustive pairwise bootstrap tests across ROIs with reliable data (with average explainable variance *>* 0.1, with more details in Methods). For every ordered region pair, we evaluated the hypothesis that one region’s switching index (*S_i_*) exceeded that of the other (*S_j_*) (null: *S_i_*≤ *S_j_*; alternative: *S_i_ > S_j_*) using bootstrap sampling across fMRI sessions. Benjamini-Hochberg false discovery rate correction[30] was applied to all resulting *p*-values. Regions that were significantly higher than others and were never significantly outperformed by any other (corrected *p <* 0.05) were defined as switching leaders (Figure 3b and Figure 4a). Thirteen such leaders emerged, distributed across the posterior and anterior bows, jointly delineating two hubs of dynamic modality reweighting across the cortex. Time resolved performance for two modalities in a switching region, Superior Frontal Gyrus, for an fMRI run is provided as an example in Figure 4c.

**Figure 3:**
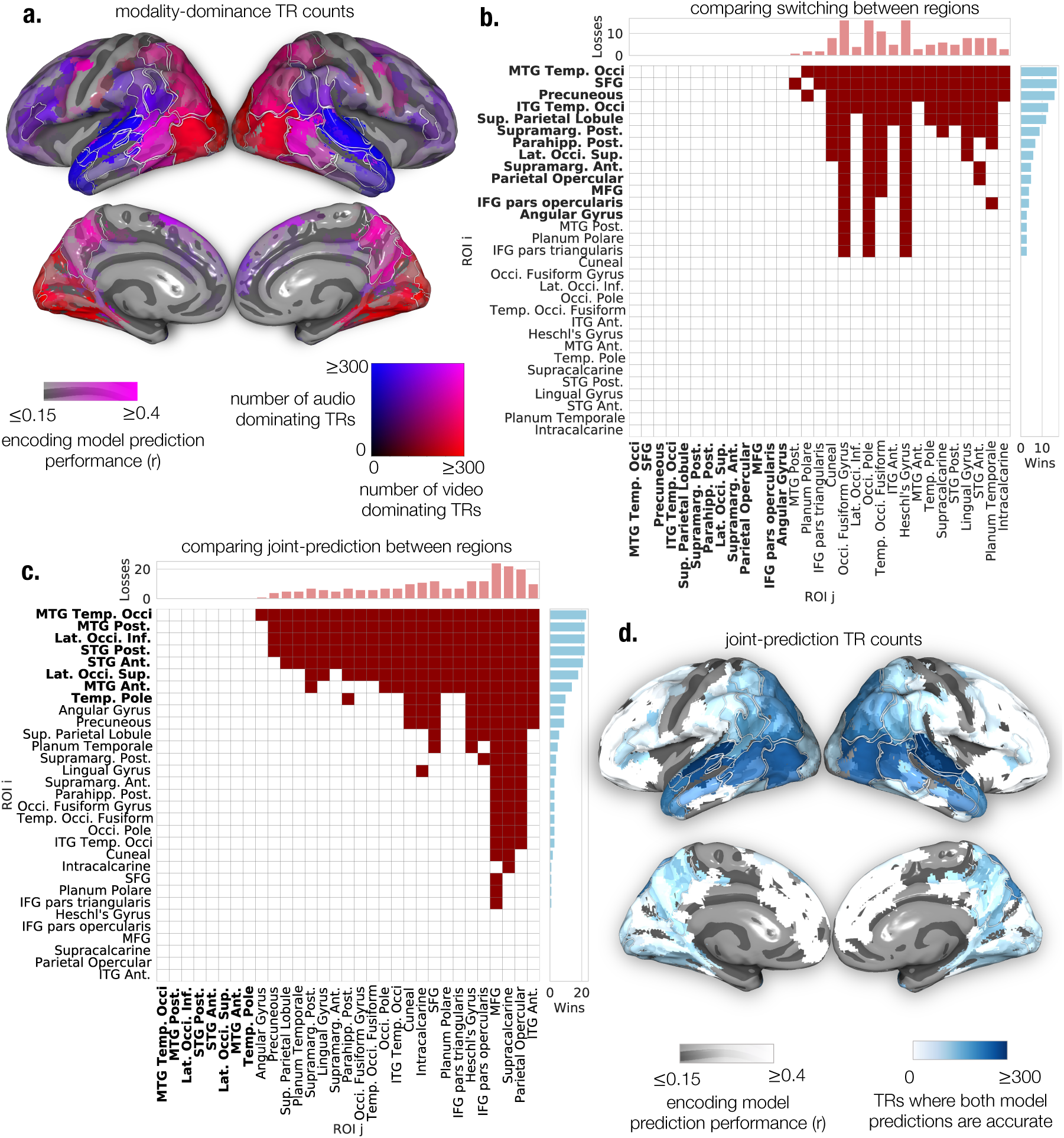
Cortical organization of modality switching and jointly-predicted dynamics. (a) Spatial distribution of the total duration of audio- and video-dominant time segments from the DIP analysis. For each cortical parcel, red intensity reflects the total duration of video-dominant segments, blue intensity reflects audio-dominant segments, and pink/purple intensity reflects parcels with substantial durations for both modalities. The encoding model prediction accuracy (Pearson r) from the concatenated audiovisual embedding model is used as a transparency layer to focus on well predicted regions. Two bands of regions, one surrounding the visual cortex and one in the frontal cortex, appear to show notable switching. (b) Results of pairwise bootstrap comparisons of the switching index. Each filled cell represents a win for region *i* (rows) over region *j* (columns), if region *i* exhibits a significantly higher switching index than region *j* (*p <* 0.05 corrected). Regions that never lose to any other region are switching leaders (shown in bold font).(c) Pairwise bootstrap comparison of the jointly-predicted index, computed analogously to (b). Each filled cell marks a win for region *i* over region *j*, meaning region *i* exhibits a significantly greater proportion of jointly-predicted timepoints after Benjamini–Hochberg correction. Bolded regions denote jointly-predicted leaders. (d) Spatial distribution of the total duration of jointly-predicted time segments from PIP. Higher blue intensity indicates parcels with longer periods where both audio and video-based model predictions are high, while the concatenated-model prediction accuracy lightens regions with low overall fit. Regions extending from the lateral occipital to the superior and middle temporal gyri are jointly-predicted leaders (shown in bold font).

**Figure 4:**
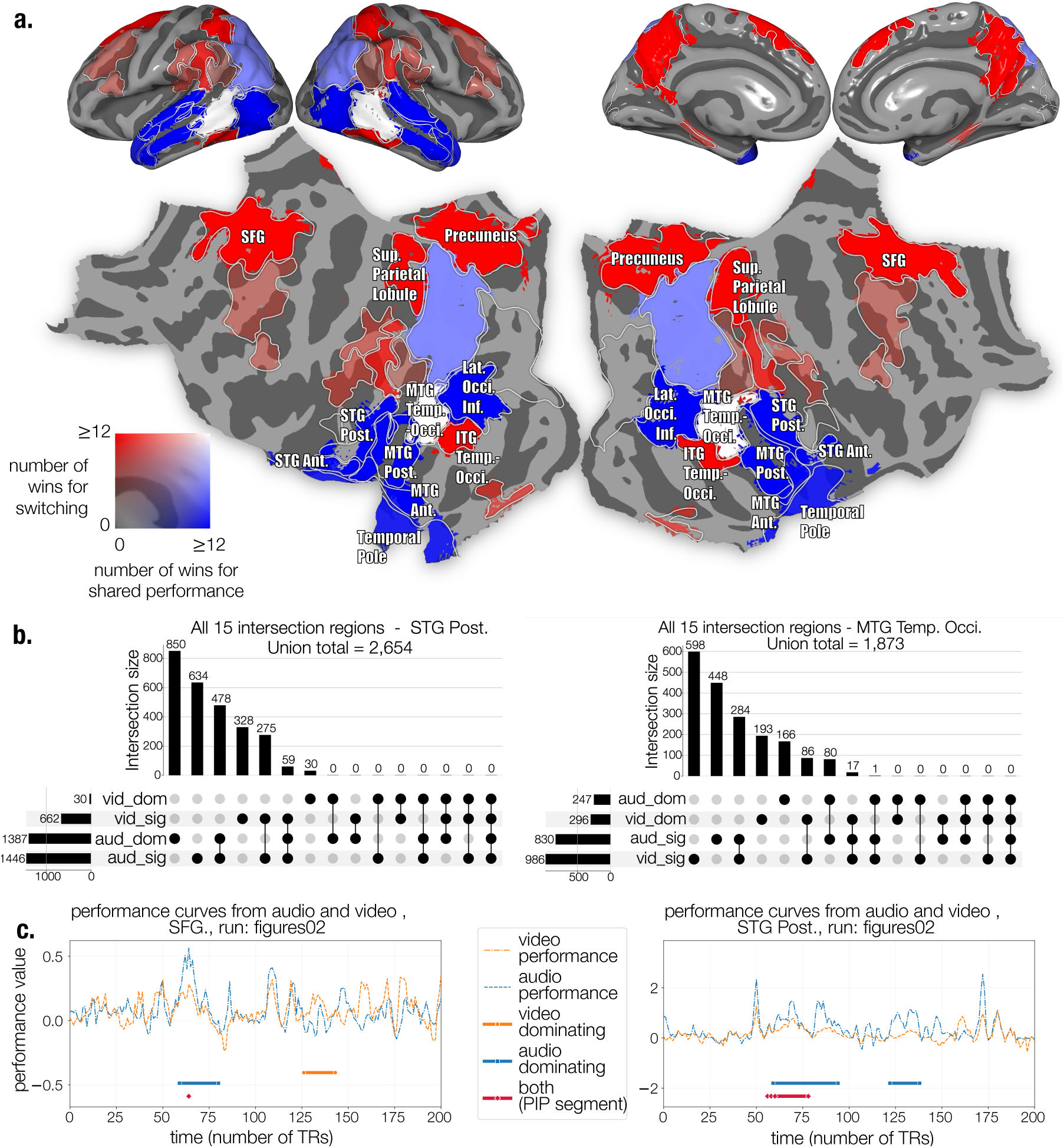
Spatial motifs and temporal trend of switching and jointly-predicted cortical regions. **(a)** Cortical distribution of *switching* and *jointly-predicted leaders*, shown by the number of significant wins each region achieved in pairwise bootstrap tests of the switching index (red) and joint-predictivity index (blue). Color intensity indicates the number of wins (see scale); regions high on both metrics appear white. Red regions form two bow-shaped structures - a posterior bow encircling visual cortex, and an anterior bow spanning lateral frontal cortex. Blue regions form an arrow-like axis extending from lateral occipital into the temporal cortex. The MTG temporo-occipital region, appearing white, is both a switching and jointlypredicted leader and lies at the intersection of the bow and arrow structures. **(b)** UpSet plots showing the size of 15 subsets of the union of 4 sets of TRs - two kinds of dominating TRs and two kinds of highly predicted TRs for two regions - STG, posterior, which is a jointly predicted leader, and MTG temporo-occipital, which is both a switching and jointly predicted leader. Most of the subsets are empty due to how these 4 sets are constrained to behave. **(c)** Examples of the per time-point performance of the audio and video model in a switching leader (SFG) and a jointly predicted leader (STG Post.) in the run called figures02 from the movie Hidden Figures. Periods identified as audio-dominant, video-dominant and predicted significantly by both modalities are marked by blue, orange and red underlines, respectively.

To assess whether periods of audio and visual dominance are consistent across regions and subjects, we ran two additional analyses. First, we asked whether a strong modality preference in one switching region coincides with the same preference elsewhere. For each pair of switching ROIs, we quantified the extent to which, when one region showed a strong modality preference as determined by DIP, the other was also significantly better predicted by that same modality than by the alternative (i.e., the performance difference in the corresponding direction was significantly above zero; Extended Figure E8). This yields a non-symmetric matrix of pairwise effect sizes, with the diagonal giving each ROI’s average preference magnitude when a preference is present. We found that 145 of 156 off-diagonal cells were significant, indicating strong overlap in preferred modality across regions; notably, 10 of the 11 non-significant cells fell in the rows or columns of parahippocampal posterior and parietal opercular cortex, suggesting potentially a more endogenous switching drive specific to these regions.

Second, we assessed consistency across subjects, using the two repeats per subject for two movies (*Hidden Figures* and *Life*) contained in the CNeuroMod Algonauts dataset. Because our main analyses averaged time courses across subjects as repetitions, these repeats allowed us instead to average the two repetitions within each subject, improving signal-to-noise ratio. We computed switching segments in a cross-validated manner, identifying segments in each held-out fold using models trained on the remaining folds and pooling segments across folds to cover both movies, and nested this procedure within a leave-one-subject-out scheme: switching segments derived from the averaged responses of three subjects were used to test whether the performance difference in the corresponding direction was significantly above zero in the held-out subject (one-sample t test, corrected for multiple comparisons), using the 13 switching regions identified in Figure 3a,b. Of the resulting 13 × 4 = 52 confidence intervals, 50 were significantly above zero (Extended Figure E7), indicating substantial cross-subject overlap. There were some region- and subject-level idiosyncrasies: subjects 2 and 5 showed somewhat less similarity than subjects 1 and 3, and ITG temporo-occipital was the most consistent region across subjects.

### An Axis of Regions spanning the occipito-temporal cortex shows remarkable joint prediction

To operationalize the performance indication framework(PIP), we again evaluated audio- and video-based encoding models on held-out movie data, this time assessing when each model independently achieved robust prediction of brain responses (Figure 2). For each ROI, we standardized each model’s performance against the permutation null at each time point, and identified TRs where the TFCE-transformed performance of each modality embedding fitted model exceeded, in at least 80% of block-wise permutations, the maximum TFCE statistic across all time points within that permutation, again controlling the familywise error rate across time points (details in Methods). We refer to these TRs as **highly predicted** by that modality and then determined the temporal overlap of highly predicted TRs of audio and visual modality. These overlapping segments marked periods when ROI responses were robustly explained by both auditory and visual features. For every ROI, we used as a metric for joint predictivity the total duration *B* of such jointly predicted segments across all sessions, yielding a measure of each ROI’s overall tendency to jointly encode auditory and visual information over time. This procedure revealed a continuous axis of jointly predicted regions extending from the lateral occipital cortex to the temporo-occipital, posterior, and anterior middle temporal gyrus, encompassing the superior temporal gyrus (anterior and posterior) and reaching the temporal pole (Figure 3d). We conducted exhaustive pairwise bootstrap tests across ROIs with average explainable variance *>* 0.1, following the same resampling procedure used for modality switching. For each region pair, we estimated the probability that one region’s joint-predictivity index (*B_i_*) exceeded that of the other (*B_j_*), and corrected all p-values using the Benjamini–Hochberg procedure. Regions that outperformed others and were never significantly outperformed by any other (corrected *p <* 0.05) were defined as jointly-predicted leaders (Figure 3c and Figure 4a).

Eight such leaders emerged, forming a continuous axis that links lateral occipital cortex to temporal cortex — a configuration that closely resembles an “arrow” of audiovisual engagement that intersects the posterior switching bow at temporo-occipital MTG (Figure 4a). Prior work has repeatedly identified the superior temporal sulcus (STS) alongside posterior STG [31, 32, 19, 33] as loci of audiovisual integration (e.g., supradditive AV responses). Although our atlas does not have a separate ROI for the STS, we observe the adjacent superior temporal gyrus parcels to be robustly predicted by both auditory and visual features, consistent with integration-related processing in nearby STS-adjacent cortex. The MTG has been implicated as another integration site using fMRI, with previous work showing that posterior STS and MTG meets the criteria for multisensory convergence, responding to both auditory and visual object stimuli and more strongly than to unimodal control stimuli [32]. A coordinate-based meta-analysis of audiovisual integration studies similarly reported convergence specifically in left MTG alongside bilateral STG [33]. Our joint-predictivity results are consistent with this: all three MTG subdivisions (anterior, posterior, and temporo-occipital) fall along the jointly-predicted axis and are among the eight joint-predictivity leaders.

To further assess how the highly predicted segments by audio or visual modality and audio and visual dominant segments interact, we use UpSet plots for all switching and jointly predicted regions, which provide a scalable alternative to Venn diagrams for visualizing overlaps among four sets. These plots show the 15 non-empty intersections defined by four sets of TRs: video-dominant (vid dom), audio-dominant (aud dom), highly predicted by video (vid sig), and highly predicted by audio (aud sig), see Figure 4b and Supplemental Figure S3. Note that audio and video dominating segments must have an empty intersection, and indeed we see aud dom ∩ vid dom = ∅, for all regions. It is possible that TRs that are video dominating are well predicted by audio, but they should be also significantly predicted by video, i.e. part of the vid dom∩aud sig∩vid sig and not the vid dom∩aud sig∩¬vid sig group (and vice versa fo the TRs that are audio dominating and well predicted by video). Consequently, many of the 15 sub-divisions are necessarily empty, given how these sets are constrained to behave. Beyond serving as a sanity check for our paradigms, this inspection reveals how STG can be strongly audio-dominant in Figure 3a while also appearing jointly predicted in Figure 3d, as the upset plots in Figure 4b reflect that these two events occur at different points in time, by quantifying TRs in each category. This is consistent with STG’s established role in auditory processing [34] alongside its involvement in audiovisual integration[32, 35] positioning it to be recruited differentially depending on the demands of the incoming stimulus..

Taken together, our results show a couple of “bow” shaped regions, one encircling the early visual cortex, and another spanning frontal regions, along with an “arrow” shaped set of regions that connect occipitotemporal regions to the temporal pole that is jointly predicted by both modalities (Figure 4a). We show an example of the time locked performances of the two modalities in a switching region (SFG) and a jointly predicted region (STG Post.) (Figure 4c). We further validate our results using a very different method-ological approach. To probe cortical reweighting more directly, we developed a multimodal attention-based model that learns content-dependent audio/video weights for each ROI at every timepoint (Figure E9a). Unlike linear models that treat each modality’s contribution as static, this architecture computes, per ROI and timepoint, a weight for audio and video that sum to 1 (a convex combination), shared across parcels within an ROI while each parcel retains its own linear readout. The model matches the performance of a combined-embedding baseline (Extended Figure E9d). For each ROI, we computed Δattention*_t_* = attention_video_(*t*) − attention_audio_(*t*) and examined its mean and standard deviation over time. Without any anatomical priors, the model recovered the brain’s canonical modality structure, assigning higher video weight to occipital regions and higher audio weight to temporal-language areas. Most switching and jointly predicted regions showed mean Δattention*_t_* near the middle of the ROI distribution (Figure E9b): switching regions showed high standard deviation, reflecting fluctuation between audioand video-weighted timepoints, while jointly predicted regions showed low standard deviation, consistent with sustained joint reliance on both modalities (Figure E9c). This convergence between modelinferred weighting and empirical dominance patterns adds more evidence in favor of the switching and jointly predicted regions.

### Dominance is not explained by low-level information or the mere presence of a preferred category

We tested whether the absence of low-level features in the non-dominant modality drives illusory switches. If dominance of one modality reflects a transient lack of input in the other, then variation in explained variance should be well predicted by variation in lowlevel features. We computed mean optical flow magnitude within each TR window as the low-level visual feature and mean spectral flux magnitude within each TR window as the low-level auditory feature, since both capture frame-to-frame change in their respective sensory content — motion and visual change for optical flow, and auditory-spectrum change for spectral flux. For each switching region, we then partitioned TRs into three classes: audio-dominant, video-dominant, and those showing neither audio nor video dominance (hereafter “neither”). Under this account, when a modality is dominant, the opposite modality should carry less lower level information than during “neither” TRs. We therefore compared the low-level auditory feature distribution of video-dominant TRs against that of “neither” TRs, and the low-level visual feature distribution of audio-dominant TRs against that of “neither” TRs. Across most regions, we found no such effect for either direction of dominance (Supplementary Figure S1). Differences in low-level feature magnitude between dominant and “neither” TRs were consistently subtle, far smaller than the corresponding differences in model performance that define dominance in the first place. Low-level feature magnitude therefore cannot account for the dominance effects we observe.

To directly compare low-level features of the non-dominant modality against how strongly that modality is encoded in the brain, we quantified both quantities across all switching regions in all three TR classes. Figure 5a contrasts “neither” TRs with video-dominant TRs (left) and with audio-dominant TRs (right), each dot representing a switching region. In both panels, the *x* coordinate gives Cliff’s delta for low-level content in the non-dominant modality and the *y* coordinate gives Cliff’s delta for variance uniquely explained by that modality. Cliff’s delta quantifies the probability that a randomly drawn observation from one group exceeds one from the other, minus the reverse, ranging from −1 to +1 [36]. Across all 13 regions and in both directions, a randomly drawn “neither” TR was more likely than a dominance TR to show higher unique explained variance in the non-dominant modality (*δ* = 0.07–0.48 for audio, 0.13–0.34 for video), and this difference exceeded the corresponding difference in low-level content (*δ* = −0.5–0.09 and −0.07–0.29, respectively), placing every region above the identity line. In several regions *δ* for low-level content was negative: dominance TRs carried *more* low-level input in the non-dominant modality than “neither” TRs, yet were less well explained by that modality’s encoding model. Dominance does not therefore appear to be driven by impoverished input, low-level features remain as rich as usual but are substantially less encoded by the brain - suggesting active sensory reweighting at those moments.

**Figure 5:**
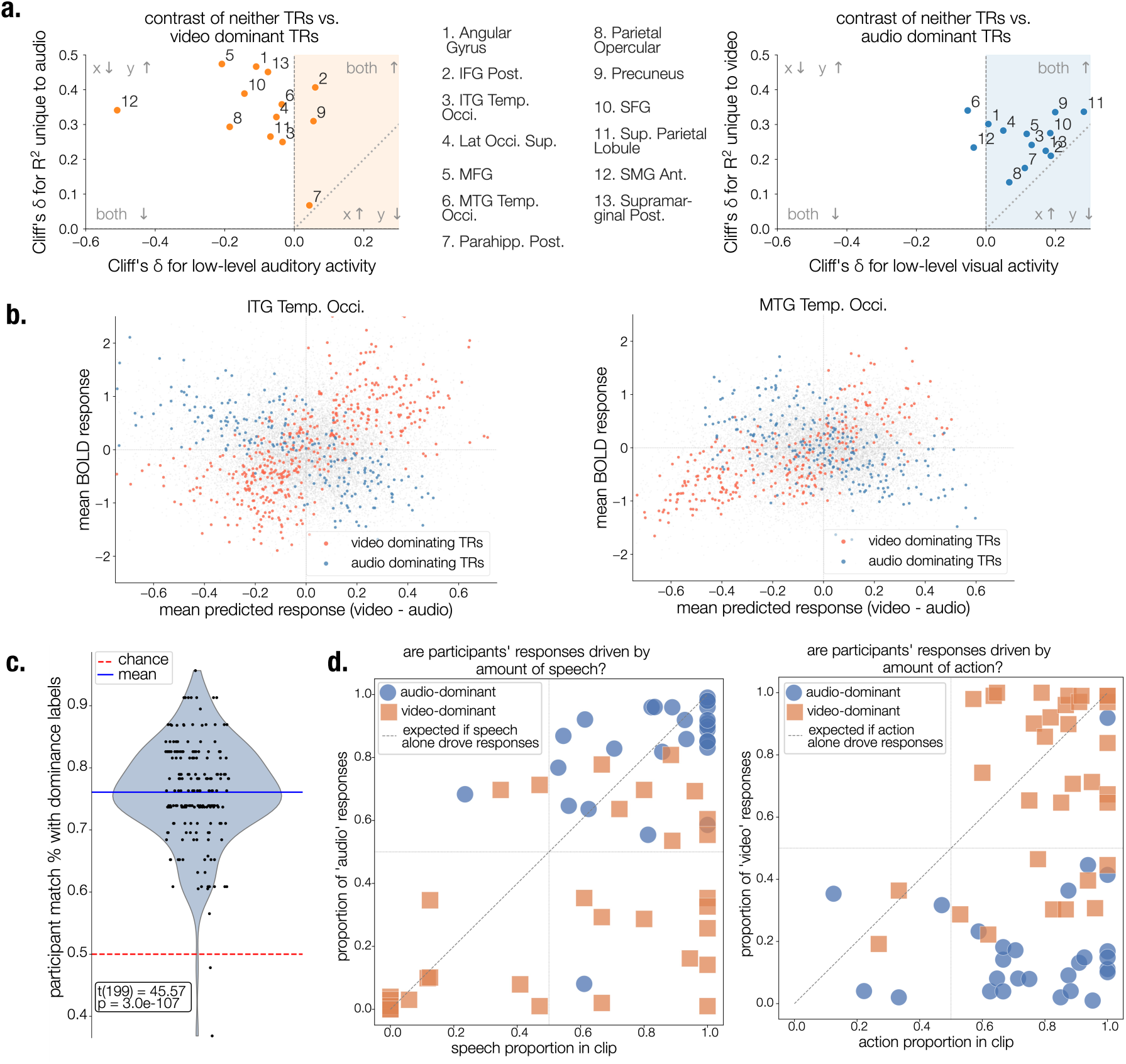
Interpretation of switching segments. **(a)** Cliff’s delta for characterizing difference between audio and video dominant TRs and usual TRs referred to as “neither” TRs. The y-axis shows the difference (neither *−* dominant) in variance uniquely explained by the non-dominant modality, and the x-axis shows the corresponding difference in low-level stimulus information. Negative x-values indicate cases where low-level information in the non-dominant modality is greater during dominance than during neither TRs. All regions lie above the diagonal, indicating that the reduction in uniquely explained variance is consistently greater than any reduction in low-level stimulus information. **(b)** Two example switching regions showing mean BOLD response (y-axis, averaged across parcels per TR) against the difference between videoand audio-model predicted responses (x-axis). Audioand video-dominant TRs are shown in blue and orange, respectively. The X-shaped pattern, observed across switching regions, argues against switching being explained solely by a region tracking its preferred concepts in whichever modality conveys them. **(c)** Participants viewed DIP-identified audioand video-dominant segments and judged which modality was more contextually important for understanding the scene. Responses matched the DIP-derived dominance labels significantly above chance (one-sample *t*-test, *t*(199) = 45.47, *p* = 3.0 *×* 10*^−^*^107^). **(d)** Relationship between participants’ modality judgments and the amount of speech or action in each clip. Points show the proportion of Audio” responses as a function of speech amount (left) and “Video” responses as a function of action amount (right), colored by DIP-derived dominance. Speech and action amount did not fully account for participants’ judgments, suggesting that they additionally reflected higher-level properties such as contextual relevance.

Several of the switching regions - including the precuneus, posterior middle and inferior temporal gyri, angular gyrus, and lateral prefrontal cortex - overlap with areas previously implicated in amodal conceptual representation[16]. We therefore asked whether switching reflects the dominant modality simply carrying the region’s preferred concepts, which could in principle be conveyed by either modality - making switching a predictable outcome of existing models under naturalistic, uncontrolled stimuli. Under this selectivity-based hypothesis, a parcel’s activity should increase whenever its preferred concept is present in either modality. Dominance segments, where one modality predicts the parcel substantially better than the other, should therefore coincide with timepoints of relatively high parcel activity due to the presence of preferred concepts.

Figure 5b shows, for two switching regions (the remainder in Supplementary Figure S2), the mean BOLD response across ROI parcels for every audio- and video-dominant TR (*y* axis) against the difference between video- and audio-predicted BOLD responses (*x* axis). The selectivity based hypothesis predicts a V-shaped pattern: under the assumption, audio-dominant TRs should fall in the negative *x* direction and video-dominant TRs in the positive *x* direction, and all the points points should fall in the positive *y* direction. Instead, all switching regions show a cross-shaped (X) pattern, with both dominance types spanning positive and negative mean BOLD responses. In many cases switching occurs when a region’s activity is relatively low, and one model correctly predicts that it should be low. Switching segments are therefore not simply moments when the concept of interest is present in one modality. Together, these analyses rule out two straightforward accounts of switching. Dominance arises neither because the non-dominant modality carries less low-level information, nor because regions simply track the presence of a preferred concept in whichever modality conveys it. What remains is an active process: the brain appears to down-weight one modality’s contribution even when its sensory input is intact and its preferred modality is not straightforwardly determined by its preferred concepts.

### Variance decomposition dissociates modality-dominant switching ROIs from integrative jointly predicted ROIs

We compared the variance components of switching and jointly predicted leader regions during audio- and video-dominated segments identified via DIP. For each ROI, we tested whether the dominant-modality unique variance (audio-unique during audio-dominated TRs; video-unique during video-dominated TRs) exceeded both the non-dominant unique component and the shared component. Four paired Wilcoxon tests were performed within each ROI (dominant-unique *>* non-dominant-unique; dominant-unique *>* shared, for both audio-dom and video-dom segments), and all p-values were FDR-corrected across all tests and ROIs (4 × N tests total).

Switching leaders exhibited significantly greater unique variance for the dominant modality (audio-unique during audio-dominated clips and video-unique during video-dominated clips) compared with both non-dominant unique and shared components (*p <* 0.05 corrected). This pattern extended across almost all switching regions, up to the lateral occipital cortex superior, whereas most jointly predicted leaders showed no statistically significant differences between dominant-modality unique and other variance components (Figure E6a). During modality-dominated segments, switching regions rely more heavily on the dominant modality’s features, while jointly predicted regions maintain more balanced modality reliance even in these segments.

Variance decomposition in jointly-predicted ROI leaders during jointly-predicted TRs revealed that shared variance was significantly higher than the audio- or video-unique components for some of the regions. During the rest of the TRs, the shared variance was not greater than the other components. One potential explanation for this result is that jointly predicted time-points correspond to moments when auditory and visual feature spaces are possibly more correlated (Figure E6b). However, this explanation alone is not sufficient, as it predicts that all highly predicted language and visual regions—including Heschl’s gyrus, occipital fusiform gyrus, occipital pole, etc would emerge as jointly-predicted leaders, which they do not. Instead, we observe that many jointly predicted regions also have a robust amount of unique variance explained by audio and video features during jointly predicted TRs, suggesting that these regions are engaged in encoding both shared and unique information from audio and video features, which would correspond to cross-modality integration or encoding of inherently multimodal constructs like emotion, social or audiovisual actions that are partially but not fully redundant across modalities.

### Behavioral judgments of contextual relevance match neural dominance labels beyond what amount of speech and action can explain

To test whether modality-dominated segments identified from switching regions reflect meaningful contextual structure in the stimuli, we conducted a behavioral experiment probing human judgments of modality importance. We first applied DIP to all switching regions treated as a single unified ROI, yielding a global set of audio-dominated and video-dominated segments. We based this decision on the results in Extended Figure E8, which shows that for a given region’s dominating segments, other regions show a significant performance difference in the same direction (e.g., during one region’s audio-dominating segments, other regions’ “audio - video” performance is also significantly above zero) and hence the switching tendency is pretty similar across regions. The participants indicated whether the audio or video was more important for understanding the central meaning of each scene and provided more contextually relevant information for what happens next in the movie. Human choices aligned closely with the DIP-derived labels: participants selected the DIP-identified dominant modality on average for 75.32% of trials, significantly above chance (*t*(199) = 45.47*, p* = 3.0*e^−^*^107^) (Figure 5c). These results suggest that the modality-dominant segments detected via DIP correspond to scenes in which one modality is contextually prioritized by human observers.

To assess whether participants’ decisions reflect low-level sensory tracking rather than instructed task demands, we compared behavioral data alignment with all switching regions against the alignment in two sensory regions, Heschl’s gyrus and occipital pole. For each region and clip, we defined the average preferred direction as “audio” if the mean “audio — video” performance of that region across TRs in that clip was positive, and “video” if negative. Each region-participant-clip combination received a binary match score, set to one if the region’s preferred direction matched the participant’s label and zero otherwise, and the matching index for a region-participant pair is the mean match score across that participant’s tested scenes. Matching indices were significantly higher in switching regions than in sensory regions (paired two-sample permutation test, BY-corrected; Extended Figure E4), indicating that participants’ identification of the plot-relevant modality draws on more than low-level sensory input.

To exclude the possibility that participants simply used the amount of speech to judge a clip as audio-dominant and the amount of dynamic action to judge it as video-dominant, rather than making a genuine judgment of contextual relevance, we quantified speech and action content in every audio- and video-dominant clip shown to participants and used these to predict participants’ response likelihood. Since movie audio tracks contain speech, nonspeech, and music, we used the TIGER model [37] to isolate the speech track, then applied basic noise filtering to obtain a binary speech presence/absence label for each TR. For action, we used Qwen2.5-VL (the same model used for our embeddings) to detect actions present in each TR via the prompt detailed in the Methods section, which explicitly excluded static actions and speaking; this yielded an analogous binary action presence/absence label for each TR. This gives us, for every clip, the proportion of TRs containing speech and the proportion containing action. The details are provided in the Methods section. Figure 5d tests whether “Audio” responses track speech content and “Video” responses track action content. Each dot is a clip, colored and shaped by DIP-assigned dominance (orange square = video-dominant, blue circle = audio-dominant); the x-axis is speech or action proportion and the y-axis is mean participant response. Two patterns emerge: participants are highly accurate, with blue points skewing toward higher audio-response proportions and orange points toward higher video-response proportions; yet many high-speech clips still correctly elicit video responses (bottom right, left panel), and many high-action clips still correctly elicit audio responses (bottom right, right panel), indicating speech/action content alone does not fully determine responses.

To test this formally, we fit two logistic regression models: one predicting, for each trial, the probability of an audio-dominant response from clip-level speech proportion, and one predicting the probability of a video-dominant response from clip-level action proportion, each with random intercepts for clip and participant to account for the non-independence of repeated trials. To determine how much of this variance was explained by relevant content proportion itself versus by clip-level idiosyncrasy, we decomposed the latent-scale variance of each model into its fixed-effect and random-intercept components, using Nakagawa and Schielzeth’s approximation for logit-link GLMMs [38]. In the audio model, speech proportion accounted for R² = 0.2914, but clip identity alone accounted for more, R² = 0.3852. In the video model, this gap was even sharper: action proportion explained almost nothing (R² = 0.0320), while clip identity accounted for the majority of the model’s explanatory power (R² = 0.6516), over 20 times the contribution of action proportion. Participant identity contributed comparably little in both models (R² = 0.0518 and 0.0506, respectively). Together, these results argue against a simple account in which dominance judgments track relevant content proportion, which would predict that content proportion explains most of the variance in both models. Instead, clip identity is the dominant source of explained variance in both cases. This suggests participants responded to something beyond raw modality content, likely a higher-level construct such as the contextual relevance of one modality over the other within a given clip, which is what participants were explicitly asked to report.

## Discussion

Our analyses uncovered two complementary forms of cortical multimodality that dynamically balance sensory evidence during naturalistic perception. Using the Dominance Indication Paradigm (DIP) and Performance Indication Paradigm (PIP), we found two circular belts of regions that alternated their dominant sensory modality over time, and a continuous occipito-temporal axis that jointly tracked both modalities, together forming a bow-and-arrow-like pattern. These findings reveal that multimodal processing is not monolithic but comprising of two interacting systems that are also spatially separated: one that flexibly switches between modalities depending on context, and another that either attends to concepts shared across modalities or integrates shared audiovisual structure to maintain perceptual coherence. Further, we show that these findings are replicated using a very different analysis paradigm that relies on modeling modality importance using the attention mechanism in a transformer.

The posterior switching bow encompasses regions including the precuneus, angular gyrus, and posterior middle and inferior temporal gyri—areas traditionally linked to amodal or conceptual representation [15, 16]. Their fluctuating dominance suggests that such territories are not just permanently supramodal but sometimes also dynamically prioritize whichever modality carries more contextually relevant or reliable information. A smaller anterior bow within the lateral prefrontal cortex exhibits the same pattern, implying that dynamic modality reweighting also extends into higher-order control regions. We present evidence that this reweighting during switching is implemented via suppression of the nondominant modality: despite carrying no less low-level information than usual, the brain encodes information from that modality substantially less, and is therefore predicted less well by it. We further show that the data does not support an account in which the switching regions are selective for some categories, and attend to the modality that carries these categories, which would imply that the dominance of a modality entirely coincides with high real and predicted BOLD activity. Instead, switching sometimes occurs when a region’s activity is relatively inhibited, with the dominant modality’s model correctly predicting that it should indeed be low, indicating a more complex process.

The spatial arrangement of the posterior bow closely parallels the organization reported in Popham *et al.* [17], which found that each category-selective patch along the anterior border of the visual cortex has an immediately anterior region selective for the same semantic category in language. The authors proposed that this border acts as a convergence zone where visual semantic information enters an amodal system through parallel, category-aligned pathways. Our results may reflect a dynamic counterpart of this motif, where some of the regions might be processing both stream of information at the same time, while others are switching between streams. An exciting hypothesis to evaluate in future work is that the switching regions can be semantically selective and prefer specific categories, however, responses to preferred categories may depend on whether those categories are conveyed by the currently prioritized stream. This explanation would be consistent with the pattern we observe of dominance frequently occurring at times when the real and the predicted BOLD responses are low. This dependence of selectivity on the importance of an information stream is also reminiscent of what is observed in binocular rivalry experiments [39].

By contrast, jointly predicted regions form a continuous occipito-temporal arrow spanning the lateral occipital gyri and the STG and MTG and reaching the temporal pole, consistent with the long-recognized role of STS–adjacent cortex and MTG in audiovisual integration [19, 31, 32]. We hypothesize that integration may partially underlie this jointly predictive organization, which can be further evaluated in future work. We also observe that regions identified as jointly predicted by both modalities do not show a single uniform signature, but at least two dissociable ones. In STG and temporal pole, only the shared variance component is elevated during jointly-predicted segments relative to baseline, while unique audio and unique video contributions remain unchanged, which seems to indicate that these regions’ might be better predicted during those segments due to the correlated structure between modalities rather than to either stream’s independent content (Extended Figure E6). In MTG and lateral occipital cortex, by contrast, unique audio, unique video, and shared variance are all more or less elevated (Extended Figure E6), suggesting these regions are instead more strongly and independently driven by both streams; which could be due to these regions’ tracking of an amodal construct (e.g., emotional, social or audiovisual action content widely present in movies) that is partially but not fully redundant across modalities. STS/STG has been strongly implicated in social interaction perception in both controlled and naturalistic paradigms [18, 40], while MTG and lateral occipital cortex have each been separately linked to emotion recognition, via face-based and scene-based routes respectively [41, 42], offering candidate explanations for why these regions might show elevated joint predictability without necessarily requiring true integrative binding of information in modalities as defined in the literature. Our frameworks also underscore the need for analytic frameworks - and possibly new terminology - suited to teasing apart the many forms audiovisual interaction can take across a continuous, content-rich naturalistic stimulus, potentially unfolding over multiple, region-specific timescales.

Our results further refine aforementioned accounts of multisensory and semantic processing. Damasio’s convergence-zone framework proposes that distributed modality-specific representations are coordinated through multiple cortical convergence sites [13]; in this sense, our jointly-predicted regions are natural candidates for convergence-related processing, whereas switching regions suggest that some such territories may dynamically prioritize one sensory stream over another. The hub-and-spoke account instead proposes a transmodal anterior temporal lobe (ATL) hub interacting with modality-specific cortical “spokes” [12]. Consistent with this view, we observe strong joint predictivity extending into the temporal pole; however, multimodal dynamics are also widespread outside ATL, including jointly-predicted occipito-temporal regions and switching temporal, parietal, and frontal regions, making a strict anatomical division between a transmodal hub and unimodal spokes difficult to accommodate. Finally, Driver and Noesselt proposed that multisensory processing arises through multiple interacting pathways, including influences from higher-order multisensory regions onto sensory-specific cortex [14]. Our findings provide a time-resolved extension of this view by dissociating regions that remain sensitive to both modalities from those that undergo sustained shifts in modality dominance. Thus, rather than contradicting these theories, our results add spatial and temporal constraints: accounts of naturalistic multimodal comprehension must explain both where sensory information converges and how its relative influence changes with contextual demands.

What allows the switching regions to determine which modality to encode? Researchers interested in the role of attention in audiovisual integration have studied the role of bottomup effects, such as stimulus saliency, or top-down effects, such as task demands and expectations [43, 14]. Such flexibility has also been formalized as changes in the relative weighting of sensory evidence, for example when attention or sensory reliability alters the contribution of auditory and visual information to multisensory inference [44]. This line of inquiry often investigates how the brain allocates gain to sensory sources (e.g. the sight and sound of the same object, or the speech of a person speaking and their mouth movements) on relatively short time scales, to increase fusion between modalities or reduce interference. Our findings are therefore not intended to introduce dynamic sensory weighting itself, but rather to characterize how such weighting unfolds during continuous naturalistic comprehension. In contrast to paradigms in which attention to a modality or feature is explicitly cued, our approach reveals modality-dominant scenes on the order of tens of seconds without instructing participants to attend to one sensory stream over the other. Indeed, work on event segmentation during narrative comprehension [45] shows that high-level regions such as precuneus and angular gyrus shift their latent event model only every tens of seconds, suggesting that these regions maintain long-duration situation models rather than rapidly changing stimulus features, though this hypothesis should be further tested in future work. While some existing work indicates that frontal regions are modulated by overt attention tasks during narratives [46], other work shows that our more posterior switching ROIs also switch their tuning towards an explicitly attended category, such as the precuneus or the posterior parahippocampal gyrus [47]. Our switching ROIs also aligns considerably with the default mode network (DMN) [48] which is classically considered to be a region that is deactivated during tasks, but has recently been shown to be involved in complex tasks that require sustained representations over multiple seconds such as narratives [7, 49], whether these tasks are internally or externally driven, as argued in De Soares *et al.* [46].

Through our behavioral experiment, we showed that switching reflects higher level contextual relevance of one modality over the other to understand the global meaning of the scene. Interestingly, the responses of the participants had a lot of agreement (see Supplemental Figure S6), with many scenes having a matching rate close to 100% between the behavioral ratings and our labels. We additionally establish that participants’ judgments of which modality is contextually relevant draws on more than low-level sensory inputs (Extended Figure E4), and in fact, even high level information such as speech and action proportions do not entirely explain participants’ modality judgments. We interpret this as evidence that participants are indeed tracking something beyond stimulus properties, likely the contextual relevance, as instructed (Figure 5d). Given that switching segments span relatively long durations, on the order of tens of seconds (Extended Figure E5), switching is more likely driven by higher-level narrative structure than by transient bursts of salient information in one modality. This points toward a top-down attentional mechanism, modulated by relevant information load across modalities. We also hypothesize that attentional resources are not always divided proportionally to the information present in the modalities, with the brain instead sometimes adopting a winner-takes-all like allocation, consistent with the suppression of the non-dominant modality we observe (Figure 5a,b). Relatedly, suppression of competing inputs has been reported in very different paradigms. Within vision, simultaneously presented stimuli suppress one another’s responses, and this suppression, although already present in V1, grows progressively stronger through V2, V4 and TEO [50]. Across modalities, when a spoken and a written story are presented concurrently, responses to the unattended stream persist in early auditory and visual cortex but are strongly suppressed in higher-order regions [51].

Several methodological and interpretive limitations warrant consideration. First, although the audio and video models used to extract embeddings have demonstrated strong performance on diverse benchmarks, we cannot fully characterize what features they encode. Their internal representations likely conflate low-level sensory features with higher-order semantics, making it difficult to pinpoint the precise information driving each region’s predictivity. Nonetheless, the models’ robust performance across speech, prosody, and visual-scene understanding tasks indicates that their embeddings capture ecologically relevant structure sufficient for brain-level mapping (as is also corroborated by the very good brain prediction performance measures our models achieve, Figure 2d). Second, spatial localization is constrained by the Harvard–Oxford cortical atlas [28], which is both probabilistic and relatively coarse: each parcel’s regional assignment reflects the probability of belonging to a region across subjects, rather than a definitive boundary, and parcel units are big and hence could sometimes exist within two or three regions sharing a boundary. As a result, ROI borders can overlap or blur fine-scale functional distinctions, particularly near the superior temporal sulcus and temporal pole. Notably, when we repeat our analyses at the parcel level, we obtain a similar general pattern of switching regions (Extended Figure E3), indicating that our results are not an artifact of the chosen atlas. Higher-resolution, subject-specific surface parcellations would further mitigate this limitation and better separate interdigitated subregions.

Switching TRs made up 1,115 of the ∼17.4k test TRs across the four movies (6.4%, ∼27.5 min) which is modest but still amounts to a substantial duration. We make three points that relate to this observed level of switching. First, the dataset is deep but narrow: it has many hours per subject but only four subjects, with no repeated presentations per subject for most of the data. A 15-TR scene with genuine modality dominance in an ROI therefore offers only ∼60 TRs (∼90 s) of data across subjects, which is limited power for detecting effects at the level of individual scenes. Aggregated across scenes, switching is reliably significant at the ROI level, so we can robustly identify switching ROIs. For individual scenes, however, we can only suggest modality dominance (note that nonetheless, the behavioural analysis does corroborate the scenes we identify). Naturalistic datasets that combine long narratives, in which switching can emerge, with many repeats or subjects are currently rare, and collecting them will be important for scene-level inference. Second, we do not necessarily expect switching to be very frequent: for most of a film, both modalities can carry relevant information and there might not be any need to switch. Third, rarity does not diminish scientific relevance. The mismatch negativity, for example, is elicited by rare auditory deviants and has nonetheless become foundational to our understanding of predictive sensory processing [52].

We also note that population-level BOLD criteria can misclassify a voxel containing only interleaved unisensory populations as multisensory [53], and some work has used fMR-adaptation to approach this dissociation directly (e.g.,[54] for face–voice integration in STS). Given our current approach, we cannot conclusively rule out the possibility that the switching phenomenon arises from a mixture of unimodal populations rather than genuine reweighting. However, the suppression effect we report in Figure 5a cannot be explained on this basis alone: the non-dominant modality does not, on average, carry much less lowlevel information, yet it is markedly less well encoded by these regions during periods of opposite-modality dominance. Thus, even if these regions comprise a mixture of unimodal populations, the population tuned to the non-dominant modality encodes substantially less information than usual during such periods, despite no substantial reduction in the information available in the stimulus itself. This pattern indicates that some form of reweighting is occurring, regardless of the underlying circuit-level architecture.

In conclusion, our findings suggest that the human brain processes long, continuous naturalistic stimuli through two complementary mechanisms: some regions switch their reliance between audition and vision as the informational content of the stimulus changes, while others are jointly driven by both modalities. Together, these mechanisms may allow the brain to extract meaning from complex, dynamic audiovisual experience.

## Methods

### Feature Collection

For both the Qwen2-Audio and Qwen2.5-VL models (open-sourced on HuggingFace, we extracted contextualized hidden-state representations using a sliding-window approach. The feature vector for the *i*-th TR was obtained by passing the stimulus segment spanning [(*i* − 5) × TR*, i* × TR] (∼9 s; 1 TR = 1.49 s) through the model (audio waveform for the audio model; silent video clip for the visual model). The window was then advanced by one TR, to [(*i* − 4) × TR, (*i* + 1) × TR], to obtain the feature vector for the (*i* + 1)- th TR, and so on. Consecutive windows therefore overlap by 5 × TR, and features are updated at every TR, matching the fMRI sampling rate without temporally downsampling the stimulus. We chose a long context window because the semantic content of naturalistic audiovisual movies unfolds over several seconds, and longer context allows the models to better capture events at this level. Consistent with this, top-performing encoding models in the Algonauts 2025 challenge, including the winning TRIBE model[55], relied on long temporal context windows and showed that context spanning multiple TRs was critical for predicting responses in higher-order regions. Within each window, the final-layer hidden states of the language model backbone were averaged across all tokens to yield a single feature vector. For the video model, stimuli were sampled at 15 frames/s.

For the video model, we used the following instruction prompt: *“You are a helpful, smart assistant. You have watched many famous movies and sitcoms throughout your training process that feature the day-to-day actions and events of the main characters. You are now watching a clip from such a movie. Carefully comprehend all the events in the video and how the scenes progress, including the important actions, emotional reactions, and changes in setting.”*

For the audio model, the prompt was: <|im start|>system\nYou are a helpful assistant.<|im end|>\n<|im start|>user\nAudio 1:<|audio bos|><|AUDIO|> <|audio eos|>\nYou have listened to many famous movies and sitcoms throughout your training process that feature the day-to-day conversations and events of the main characters. You are now listening to such a clip from a movie. Carefully listen to what events are happening in the audio, what the characters are talking about, and their intentions.<|im end|>\n<|im start|> assistant\n

Because each feature vector summarizes the preceding 6 × TR of stimulus, the first complete window is available only at the 6th TR. Accordingly, audiovisual features were aligned to the fMRI data from that point onward, and only those volumes were used for model fitting and testing.

For VideoMAE feature extraction, each 4 × TR segment was uniformly sampled at 16 frames, which were passed through the pretrained model. We collected the hidden states from the final transformer layer and averaged them across the sampled frames. The first two feature segments were discarded to maintain temporal alignment with the other feature sets.

### Explainable Variance Computation

To estimate the reliability of fMRI responses across subjects and thereby quantify the explainable variance of each parcel, we computed a noise-ceiling measure based on repeated presentations of the same stimuli [56]. For each fMRI session, data from four participants viewing identical segments were treated as repeated measurements. Let *X* ∈ R*^R×T^ ^×V^* denote the data matrix, where *R* is the number of repeats (subjects), *T* the number of time points, and *V* the number of parcels or voxels. Each time course was *z*-scored along the temporal dimension to remove baseline differences across repeats. For each parcel *v*, the residual variance across repeats was computed as

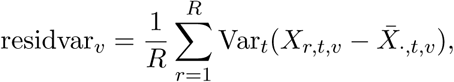

and the explainable variance was then defined as

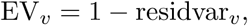

corresponding to the proportion of response variance that is reproducible across subjects. A small-sample bias correction was applied based on the number of repeats,

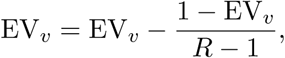

following the same formulation used in prior noise-ceiling analyses. This measure served to identify parcels with a sufficiently reliable signal for downstream analyses.

### Dataset and Encoding model fitting

The Algonauts Project-2025 challenge[23] aims to predict human brain responses to complex, naturalistic audiovisual stimuli. Using the extensive dataset collected by the Courtois Project on Neuronal Modeling (CNeuroMod) team [22], it provides the largest publicly available fMRI dataset for modeling neural responses to multimodal movies. We used a subset of the released *FRIENDS* data, which includes the first three seasons, and the four available full-length films—*The Bourne Supremacy*, *The Wolf of Wall Street*, *Hidden Fig-ures*, and the documentary *Life*. The total duration of fMRI data we used was more than 36 hours. While *FRIENDS* is relatively dialogue-heavy and therefore biased toward auditory content, the movies offer a more balanced mix of visual and auditory information: *Bourne Supremacy* and *The Wolf of Wall Street* feature rich visual dynamics, whereas *Hidden Figures* and *Life* emphasize speech, social interaction, and narrative structure.

The fMRI data were acquired from English-speaking participants recorded in a 3T Siemens PrismaFit scanner. Four participants’ data was made available in the Algonauts competition. Each Friends episode and movie was divided into fMRI runs of about 12 minutes. Participants watched segments sequentially, though not necessarily in a single session. The data was preprocessed by the Courtois Neuromod group using fMRIPrep [57], which included motion correction, slice-timing correction, and co-registration of the functional images of each participant to their T1-weighted anatomical scan. The fMRI time series were resampled to the MNI152NLin2009cAsym and then averaged into 1000 parcels from the Schaeffer atlas [24]. For more information on data preprocessing, please see the Courtois Neuromod documentation website [58]. We further averaged the parcel data across participants.

To ensure a fair evaluation of modality-specific and multimodal effects, we adopted a two-split testing scheme. For each movie, half of the fMRI sessions were included in the training set together with the *FRIENDS* data, and the remaining sessions were held out for testing and we covered all movies this way, which consists of about 17.4k total TRs. All reported results are based on these held-out movie sessions.

To fit the encoding models, we used the extracted audiovisual features to predict parcelwise fMRI responses. To account for the hemodynamic lag, we concatenated features from the current TR and the preceding five TRs when predicting the BOLD response at the current time point. The concatenated feature vectors were reduced to 1000 dimensions via principal component analysis (PCA). For audio features, all components were taken from a single model; for video features, we combined two models by reducing each to 500 dimensions and concatenating them to match the 1000-dimensional audio representation. Separate ridge regression models were then trained for the audio and visual features on the training set, and model performance was evaluated on the held-out movie sessions as described above, by computing Pearson correlation across time.

To define our ROIs, we use the Harvard–Oxford cortical atlas [28]. We further compute another performance measure for evaluating the time dynamics of encoding model predictions. We take the dot product of the data and the predicted data across the parcels in an ROI for each time point (see Figure 2c) and divide the product by the number of parcels in that ROI. This yields a measure of model fit for each ROI and each time point.

### Common procedure for the Dominance and Performance Indication Paradigms

DIP and PIP share a similar statistical procedure, differing primarily in the two time-resolved vectors to which it is applied. In both cases, for each ROI and each fMRI run (around 12 minutes), we obtain a pair of per-time-point vectors: the two directional performance-difference series for DIP, and the audio and video model performance series for PIP. Each vector is treated independently by the steps below.

Null distributions were generated by randomly shuffling the measured fMRI responses in contiguous blocks of seven TRs, with the encoding models and their predictions held fixed, and recomputing model performance against each shuffled response series. This blockwise permutation preserves the temporal autocorrelation of the BOLD signal while disrupting its true alignment with the model predictions. Because a single shuffled response series is evaluated against both the audio and video models, any shuffle-induced variance common to the two is shared rather than contributing spuriously to their comparison.

Because chance-level model performance varies across time points, we standardized the value at each time point *t* using the mean *µ_t_* and standard deviation *σ_t_*of the pooled set comprising the observed and all permuted values at that point, placing all time points on a common scale before cluster enhancement. The identical transform is applied to the observed and to every permuted vector, so exchangeability is preserved and the test remains exact. This is the non-parametric analogue of applying TFCE to a *t*-statistic image rather than a raw contrast image [29, 59, 60].

TFCE was then applied to each standardized vector over the same range of thresholds *h* ∈ [0.01, 0.3] in increments of Δ*h* = 0.01,

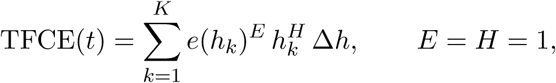

where *e*(*h_k_*) denotes the temporal extent, that is, the number of consecutive time points for which the standardized value exceeds threshold *h_k_*, and *E* and *H* are the extent and height exponents. This integral-like operation enhances contiguous segments of strong values while penalizing isolated fluctuations.

Significance was assessed against the maximum-statistic null. For each permutation we retained the maximum TFCE score across all time points, and a TR at position *p* was labeled significant when

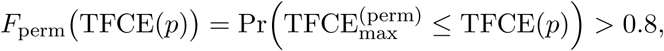

where TFCE(*p*) is the observed TFCE value at *p*, TFCE^(perm)^ is the maximum TFCE value under permutation, and *F*_perm_(·) its cumulative distribution function. Comparing against the permutation-wise maximum rather than a time-point-specific null controls the family-wise error rate across time points [59, 29].

### Dominance Indication Paradigm (DIP)

To quantify modality dominance, we computed for each ROI the per-time-point difference in model fit between the video-based and audio-based encoding models, yielding two directional performance-difference series: one capturing where the video model outperformed the audio model, and the other the reverse. Applying the procedure above to each series labels a TR as **audio-dominant** (for the audio − video direction) or **video-dominant** (for the reverse). These dominant TRs represent moments when one modality uniquely accounts for more variance in the neural signal than the other, beyond chance fluctuations estimated by the permutation baseline.

We collected dominating TRs from both modalities for each ROI across every fMRI run. For each ROI, we computed the total duration of audio-dominant segments (*D*_audio_) and of video-dominant segments (*D*_video_), and defined the switching index as *S* = min(*D*_audio_*, D*_video_). This identifies ROIs that switch multiple times and switch to both modalities.

### Performance Indication Paradigm (PIP)

PIP applies the same procedure directly to the time-resolved prediction performance of each modality rather than to their difference. For each ROI and run, the audio and video model performance vectors were standardized and TFCE-transformed independently, and TRs exceeding the significance criterion were labeled **highly predicted** for that modality.

This yields, as its primary output, the set of TRs at which each modality alone predicts the ROI well. In a post-hoc step, we intersected the highly predicted TRs from the audio and video analyses within each run to obtain the set of **jointly-predicted TRs**, corresponding to time points where both modalities robustly explained variance in the neural response. For each ROI, the total length of jointly-predicted TRs, *B*, serves as the joint-prediction index.

### Pairwise bootstrap testing

We use a two-level statistical framework: first, block-permutation TFCE testing identifies segments within each region where audio-dominance, video-dominance, or joint prediction exceeds chance across time. Second, since no analytic null exists for switching/joint-prediction tendency of a region, we define explicit metrics and compare them pairwise across regions to isolate those with disproportionately high switching or joint-prediction propensity. Pairwise, one-sided bootstrap tests were used to assess whether the switching index (*S*) or joint-predictivity index (*B*) of one region exceeded that of another. For each ordered region pair (*i, j*), the null hypothesis (*H*_0_ : *M_i_*≤ *M_j_*) was evaluated separately for *M* ∈ {*S, B*}. In each test, the indices *M* from individual fMRI runs were resampled with replacement 20,000 times. For each bootstrap sample, the sum of the resampled indices was computed for each ROI and compared between pairs of ROIs. This procedure yielded an empirical sampling distribution of pairwise differences (*M_i_* − *M_j_*), from which the one-sided *p*-value was estimated as the proportion of iterations where *M_i_* − *M_j_* ≤ 0 (alternative: *M_i_ > M_j_*). Exhaustive pairwise comparisons were conducted among all regions with mean explainable variance above 0.1, and all resulting *p*-values were corrected for multiple comparisons using the Benjamini–Hochberg false discovery rate[30] procedure (*p*_corr_ *<* 0.05). Regions that were never significantly outperformed by any other were designated as “leaders” for the corresponding measure (*S* for switching, *B* for joint predictivity).

### Behavioral Experiment

We ran a behavioral experiment approved by the Carnegie Mellon University Institutional Review Board. The switching clips total 28 min of video, but participants viewed them one at a time and made a decision after each clip, so the full set would have required substantially longer to go through. To limit fatigue, we split the stimulus set into two groups, each viewed by ∼100 participants recruited via Prolific. DIP was applied to the switching network considered as a single unified ROI to derive a global set of audio- and video-dominated segments. On the welcome screen, the following instruction was provided - In this experiment, you’ll watch short clips (average length 20s) from four Hollywood movies. Your task is to decide whether the audio (speech and non-speech sounds) or the video (visual content) is more important for understanding the core meaning of the scene. A helpful way to think about this is to ask yourself: Which modality is providing information that feels especially relevant for what happens next in the movie? We will show a few examples before the actual trials to make the idea clearer. This study usually takes around 25 minutes.

Before the main task, participants completed four instructional examples demonstrating how contextual relevance can make one modality more informative than the other to help them understand the task better. On each trial, participants viewed one of the DIP-identified segments and judged which modality—audio or video—was more important for understanding the core meaning of the scene. Responses were collected via a two-alternative multiple-choice question presented immediately after the clip ended.

### Speech and action amount characterization

For computing amount of action in every clip, we used the model Qwen2.5VL-7B-instruct that we have also used the embeddings from. We collect the actions present in every TR using the following prompt You are watching a short video clip. List every distinct physical action performed by any character or object in this clip, excluding speaking, talking. Also exclude static states or postures with no movement, such as standing, sitting, holding an object, or lying down We next construct a binary indicator for every TR using the model generated json files that indicates presence/absence of actions and thus quantify the proportion of action in every switching segment, which typically encompasses 10s of TRs.

Speech labels for each TR were derived from the speech audio stem isolated using the TIGER model [37]. The speech waveform was rectified and low-pass filtered (4th-order Butterworth, zero-phase) with a cutoff corresponding to a 200 ms smoothing window, bridging brief inter-word pauses so they were not misclassified as silence. The resulting envelope was normalized to the peak amplitude of each episode and segmented into non-overlapping 1.49 s windows matching the fMRI TR. For each window, the mean envelope amplitude was converted to decibels relative to the episode’s peak, and a binary speech-presence label was assigned (1 if the window’s mean amplitude exceeded -35 dB, 0 otherwise)

We then fit two simple mixed effect logistic regression models using the following two equations

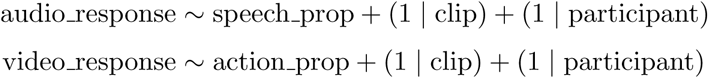

Finally, to formally quantify how much of the total variance in response probability was attributable to each predictor alone versus the full model including random effects, marginal and conditional R² were computed using the Nakagawa and Schielzeth latent-scale approximation for logit-link GLMMs, which decomposes total variance into variance explained by fixed effects, variance attributable to each random intercept, and a fixed distribution-specific term for the logit link.

### Low-level audiovisual features

To index moment-to-moment low-level sensory change in each modality, we computed spectral flux (audio) and optical flow magnitude (video) for each movie run and averaged each measure within TR-length windows (TR = 1.49 s). This gave one value per TR, aligned to the fMRI acquisition.

#### Spectral flux

We extracted the audio track with FFmpeg, downmixed it to mono and resampled it to 22,050 Hz. We then computed the magnitude short-time Fourier transform (librosa; 2,048-sample Hann window, ∼92.9 ms; hop length 512 samples, ∼23.2 ms). Spectral flux at frame *n* was defined as the L2 norm of the half-wave-rectified spectral difference,

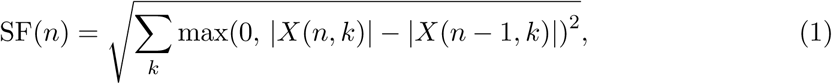

where |*X*(*n, k*)| is the STFT magnitude at frame *n* and frequency bin *k*. Only increases in spectral energy (onsets) therefore contributed. Frame-wise values were assigned to TRs by timestamp and averaged within each TR.

#### Optical flow magnitude

Video frames were converted to greyscale and downsampled to half resolution. We estimated dense optical flow between consecutive frames using the Farneback algorithm (OpenCV; pyramid scale 0.5, 3 pyramid levels, 15-pixel averaging window, 3 iterations, 5-pixel polynomial expansion neighbourhood, *σ* = 1.2). For each frame pair, we took the mean flow-vector magnitude across pixels and assigned it to the timestamp of the later frame. These values were then averaged within each TR.

### Transformer training and evaluation

To capture how each cortical region dynamically attends to auditory and visual information over time, we designed a transformer-based model that learns modality-specific attention weights for every region that vary across time. At each time point, audio and video embedding vectors (*x_a_, x_v_* ∈ R*^B×^*^1^*^×^*^1000^) were independently processed by audio and video transformer encoders, each comprising two layers (*N_a_* = *N_v_* = 2) of multi-head self-attention with four attention heads (*n*_heads_ = 4) and feedforward sublayers of dimension 256 (*d*_ff_ = 256), each followed by pre-layer normalization and dropout (*p*_dropout_ = 0.5). Learned positional embeddings (pos*_a_,* pos*_v_*) were added to preserve order, after which the encoded representations were concatenated along the modality dimension to yield *h* = [*h_a_*; *h_v_*] ∈ R*^B×^*^2^*^×^*^1000^. Region-of-interest (ROI) level query matrices *Q*_roi_ ∈ R*^N^*^roi^*^×^*^1000^ were used to compute attention weights over the two modalities via

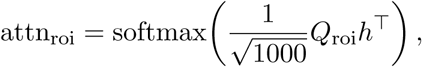

producing **ROI-specific attention distributions that were broadcast to all parcels within each ROI** according to parcel-ROI assignments. For each parcel, a modalityweighted context vector was then computed as

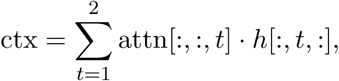

followed by dropout (*p*_context dropout_ = 0.3) to prevent overfitting. Parcelwise BOLD predictions were generated through independently learned linear readouts with parameters *W*_read_ ∈ R*^N^*^parcel^*^×^*^1000^ and *b* ∈ R*^N^*^parcel^, optimized jointly with all other model parameters during training. The model was trained to minimize mean squared error between predicted and observed parcel responses using Adam (learning rate 10*^−^*^4^, batch size 400, weight decay 10*^−^*^2^), with early stopping if validation loss failed to improve for 15 consecutive epochs. For *inference* on held-out stimuli, we passed the audio/video embeddings through the trained encoders, computed ROI-level attention weights from the learned query matrix, broadcast them to parcels, and formed a parcel-level representation *z* ∈ R*^B×N^*^parcel^*^×^*^1000^ by the attention-weighted sum over the two modality embeddings. Predictions were then obtained as

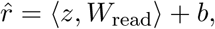

using the trained *W*_read_ and *b*. The audio and video attention weights used to compute the predicted responses were taken from this same held-out forward pass and used for the analysis reported in Figure E9.

## Supporting information

Supplemental material referred to in the manuscript text

## Acknowledgements

This research was supported by NSF CAREER grant 2237064. The authors thank the Courtois Neuromod team for collecting the large amount of fMRI data and the Algonauts team for creating a platform and a competition to make it available. The authors also thank Michael Tarr, Jenelle Feather and Bradford Mahon for scientific discussions of the work, Aaditya Ramdas for insightful discussions regarding the statistical analyses, and Isil Bilgin for assistance with the dataset.

## Extended figures

**Extended Figure E1:**
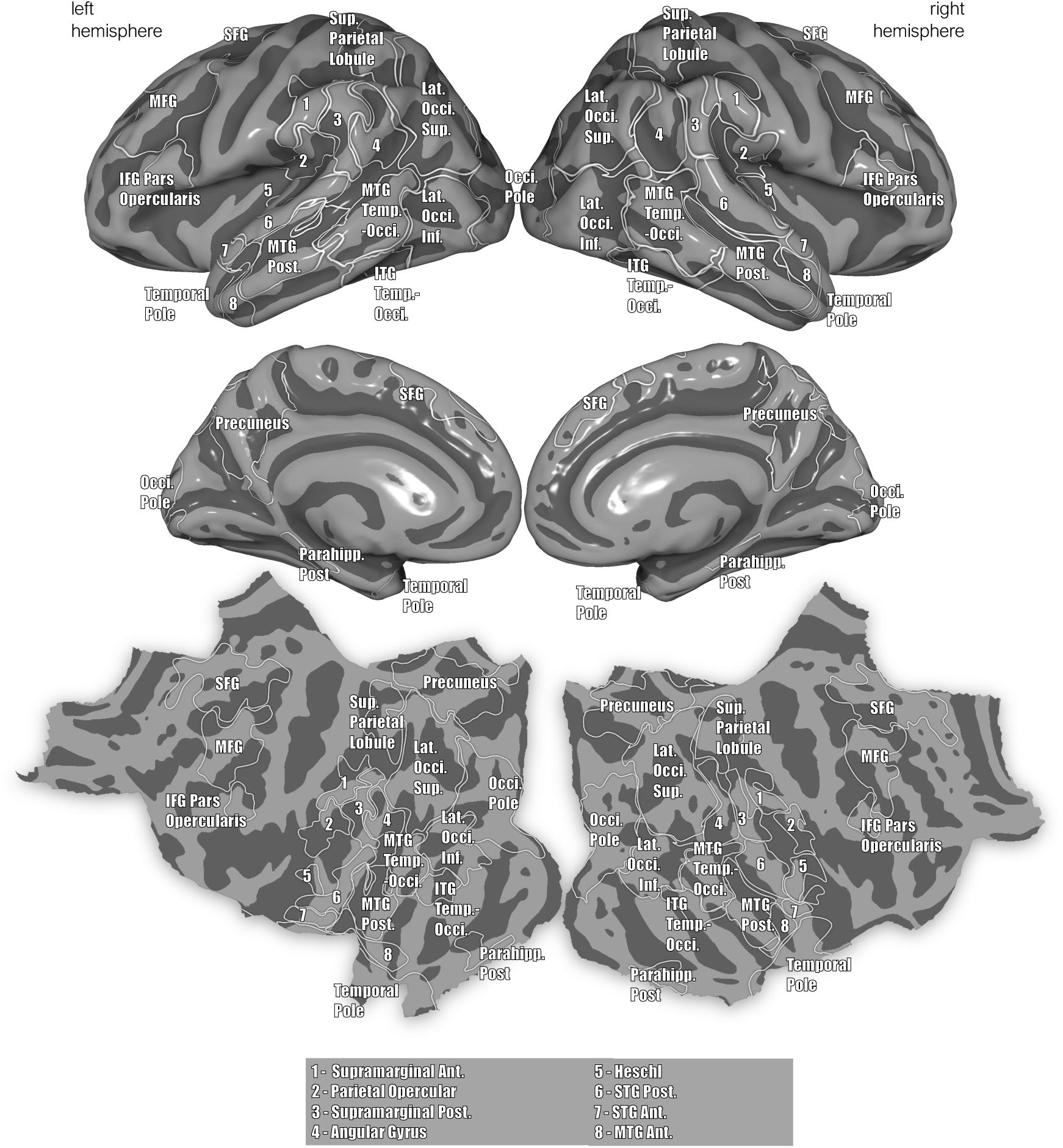
Labeled ROIs with notable prediction performance on the inflated and flattened surface. Abbreviations as follows. MFG: middle frontal gyrus, SFG: superior frontal gyrus, IFG: inferior frontal gyrus, MTG: middle temporal gyrus, STG: superior temporal gyrus, ITG: inferior temporal gyrus, Sup.: superior, Inf.: inferior, Lat.: lateral, Ant.: anterior, Post.:posterior, Occi.: occipital, Temp.: temporal, Parahipp.: parahippocampal.

**Extended Figure E2:**
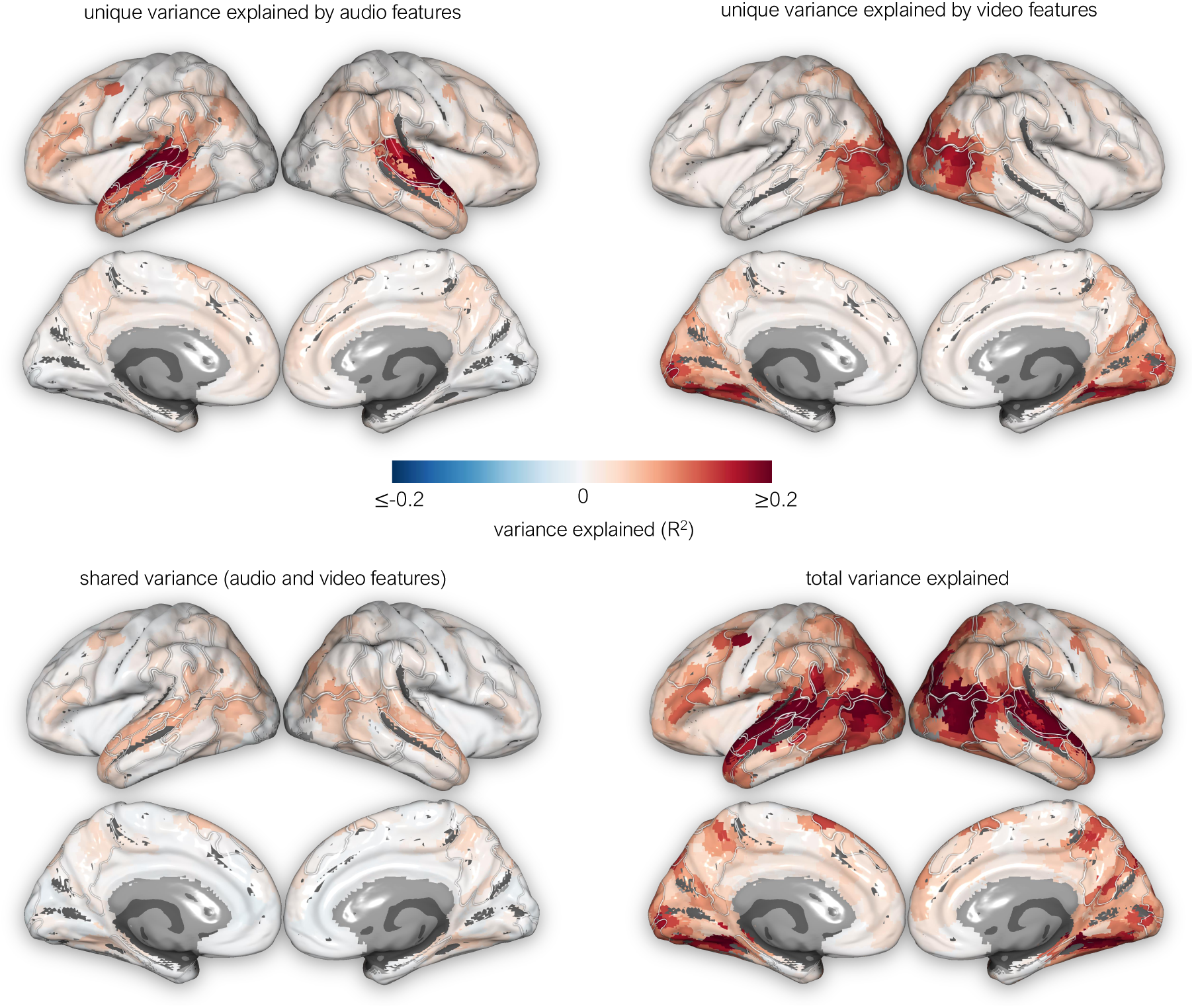
Components of explained variance, showing variance uniquely explained by the audio (top left), uniquely explained by the video (top right), jointly explained by both modalities (bottom left), and the total variance explained (bottom right).

**Extended Figure E3:**
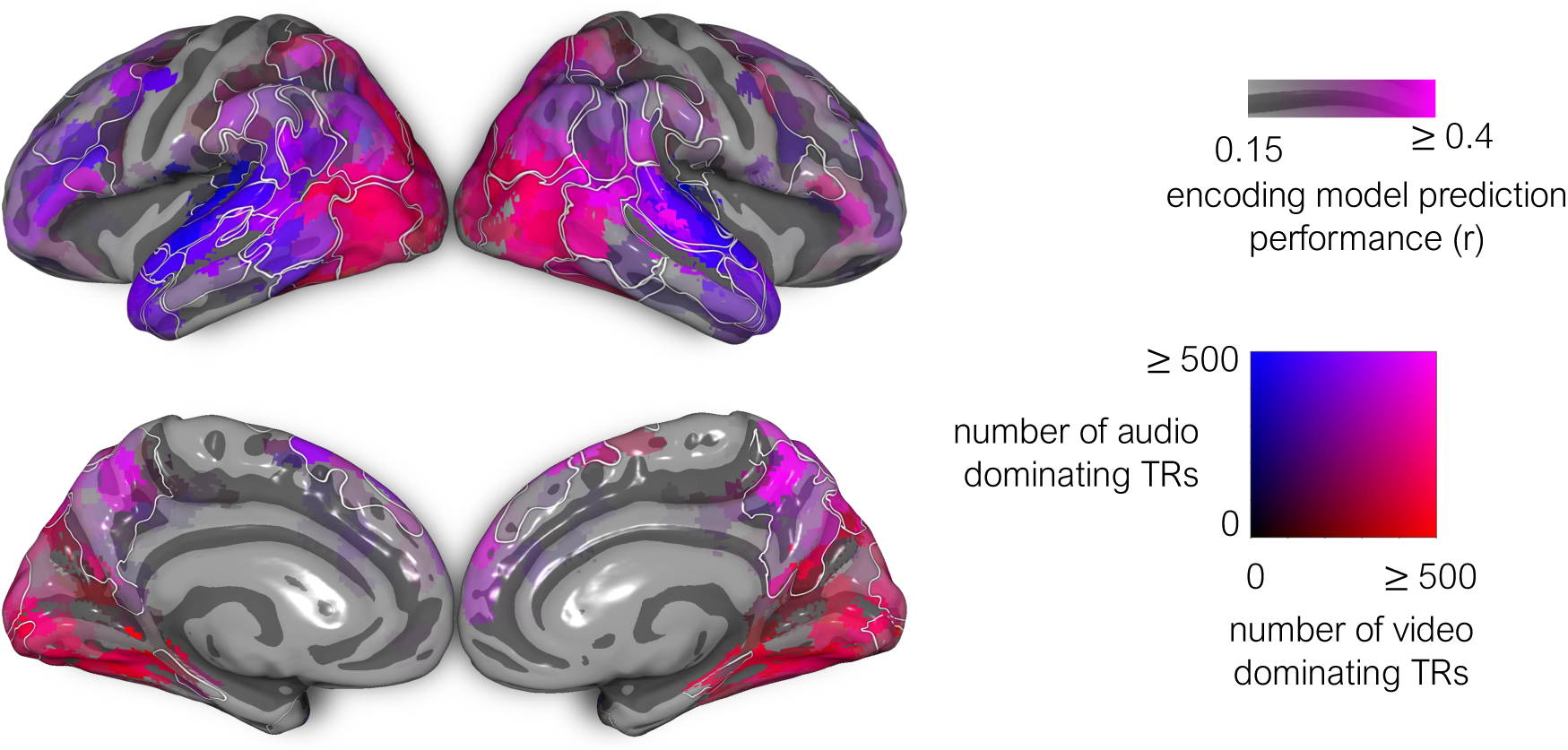
Each parcel is colored according to the total duration of time segments across all episodes where one modality’s prediction performance exceeded the other’s by *≥* 0.05 for at least 10 consecutive TRs. Redness of a parcel indicates total duration of segments where video-based predictions surpassed audio-based predictions that way; blueness of a parcel indicates the reverse. Thus parcels appearing pink or purple mean they exhibit both trends—reflecting frequent reversals in predicted dominance. Regions identified as “switching” via formal DIP analysis tend to include a mixture of such pink and purple parcels, along with pockets of predominantly audio- or video-biased parcels.

**Extended Table E1:**
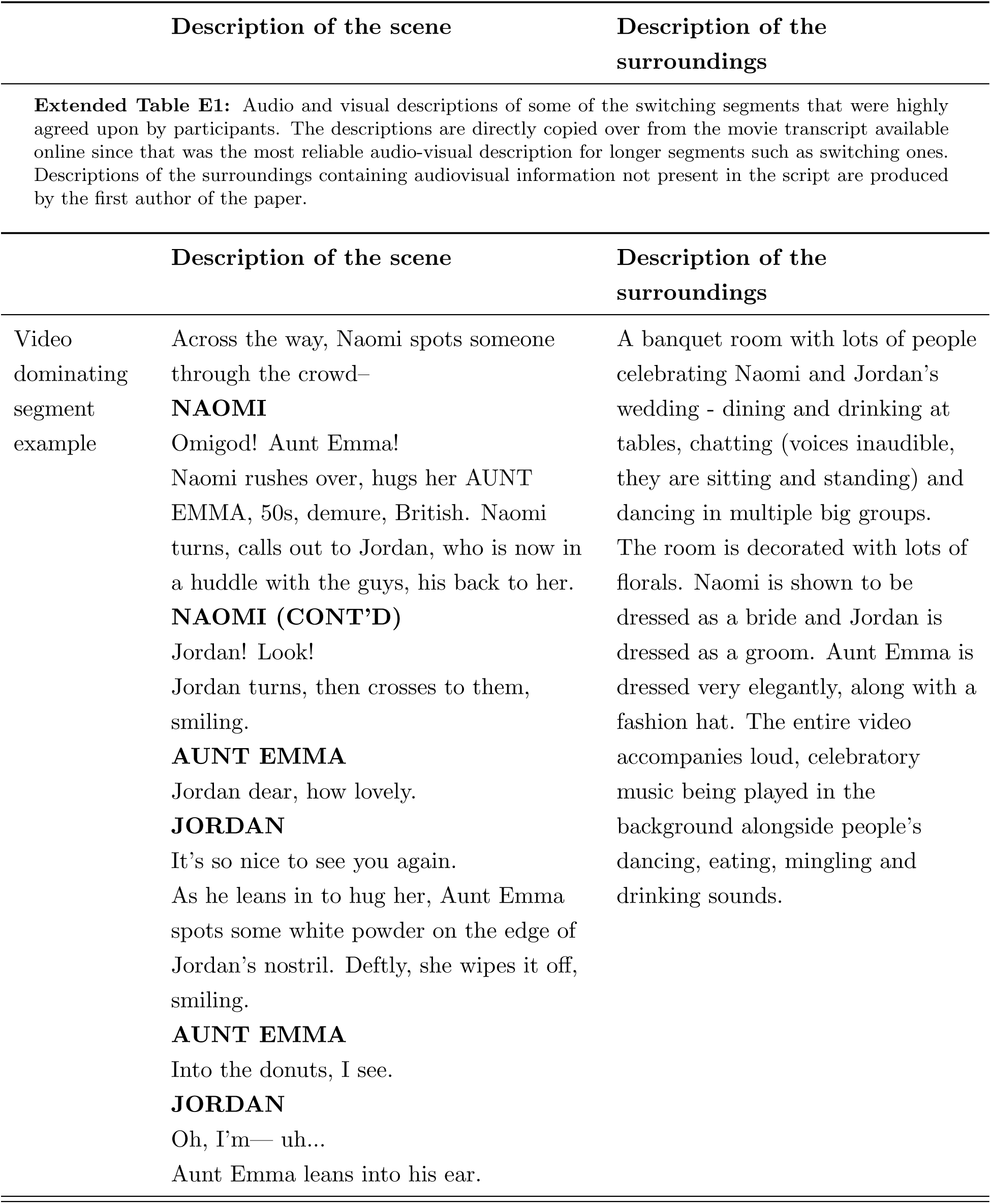
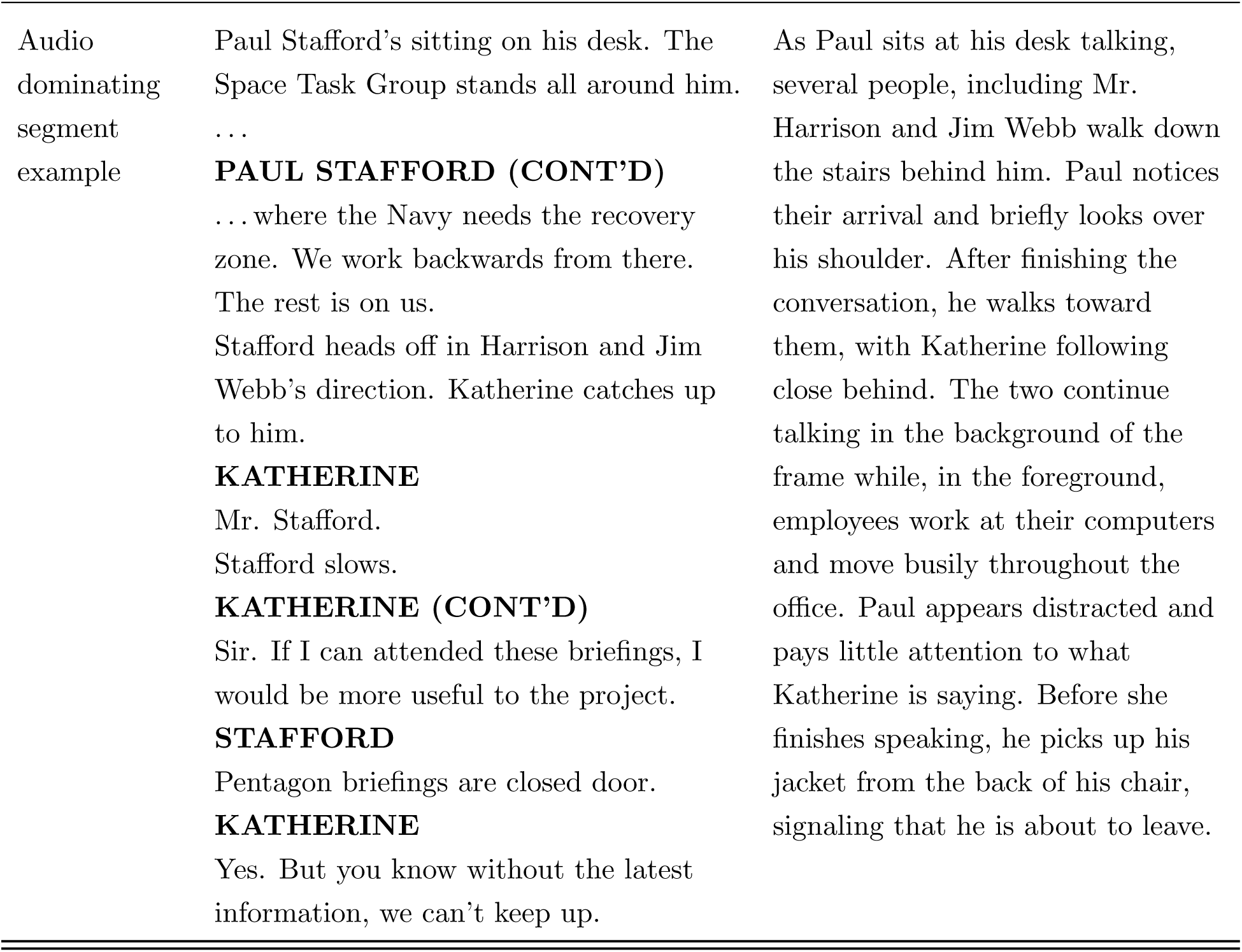
Audio and visual descriptions of some of the switching segments that were highly agreed upon by participants. The descriptions are directly copied over from the movie transcript available online since that was the most reliable audio-visual description for longer segments such as switching ones. Descriptions of the surroundings containing audiovisual information not present in the script are produced by the first author of the paper.

|  | Description of the scene | Description of the surroundings |
| --- | --- | --- |
|  | Description of the scene | Description of the surroundings |
| Video dominating segment example | Across the way, Naomi spots someone through the crowd—<br>**NAOMI**<br>Omigod! Aunt Emma!<br>Naomi rushes over, hugs her **AUNT EMMA**, 50s, demure, British. Naomi turns, calls out to Jordan, who is now in a huddle with the guys, his back to her.<br>**NAOMI (CONT'D)**<br>Jordan! Look!<br>Jordan turns, then crosses to them, smiling.<br>**AUNT EMMA**<br>Jordan dear, how lovely.<br>**JORDAN**<br>It's so nice to see you again.<br>As he leans in to hug her, Aunt Emma spots some white powder on the edge of Jordan's nostril. Deftly, she wipes it off, smiling.<br>**AUNT EMMA**<br>Into the donuts, I see.<br>**JORDAN**<br>Oh, I'm— uh...<br>Aunt Emma leans into his ear. | A banquet room with lots of people celebrating Naomi and Jordan's wedding - dining and drinking at tables, chatting (voices inaudible, they are sitting and standing) and dancing in multiple big groups.<br>The room is decorated with lots of florals. Naomi is shown to be dressed as a bride and Jordan is dressed as a groom. Aunt Emma is dressed very elegantly, along with a fashion hat. The entire video accompanies loud, celebratory music being played in the background alongside people's dancing, eating, mingling and drinking sounds. |
| Audio dominating segment example | Paul Stafford's sitting on his desk. The Space Task Group stands all around him. ...<br>**PAUL STAFFORD (CONT'D)**<br>... where the Navy needs the recovery zone. We work backwards from there. The rest is on us.<br>Stafford heads off in Harrison and Jim Webb's direction. Katherine catches up to him.<br>**KATHERINE**<br>Mr. Stafford.<br>Stafford slows.<br>**KATHERINE (CONT'D)**<br>Sir. If I can attended these briefings, I would be more useful to the project.<br>**STAFFORD**<br>Pentagon briefings are closed door.<br>**KATHERINE**<br>Yes. But you know without the latest information, we can't keep up. | As Paul sits at his desk talking, several people, including Mr. Harrison and Jim Webb walk down the stairs behind him. Paul notices their arrival and briefly looks over his shoulder. After finishing the conversation, he walks toward them, with Katherine following close behind. The two continue talking in the background of the frame while, in the foreground, employees work at their computers and move busily throughout the office. Paul appears distracted and pays little attention to what Katherine is saying. Before she finishes speaking, he picks up his jacket from the back of his chair, signaling that he is about to leave. |

**Extended Figure E4:**
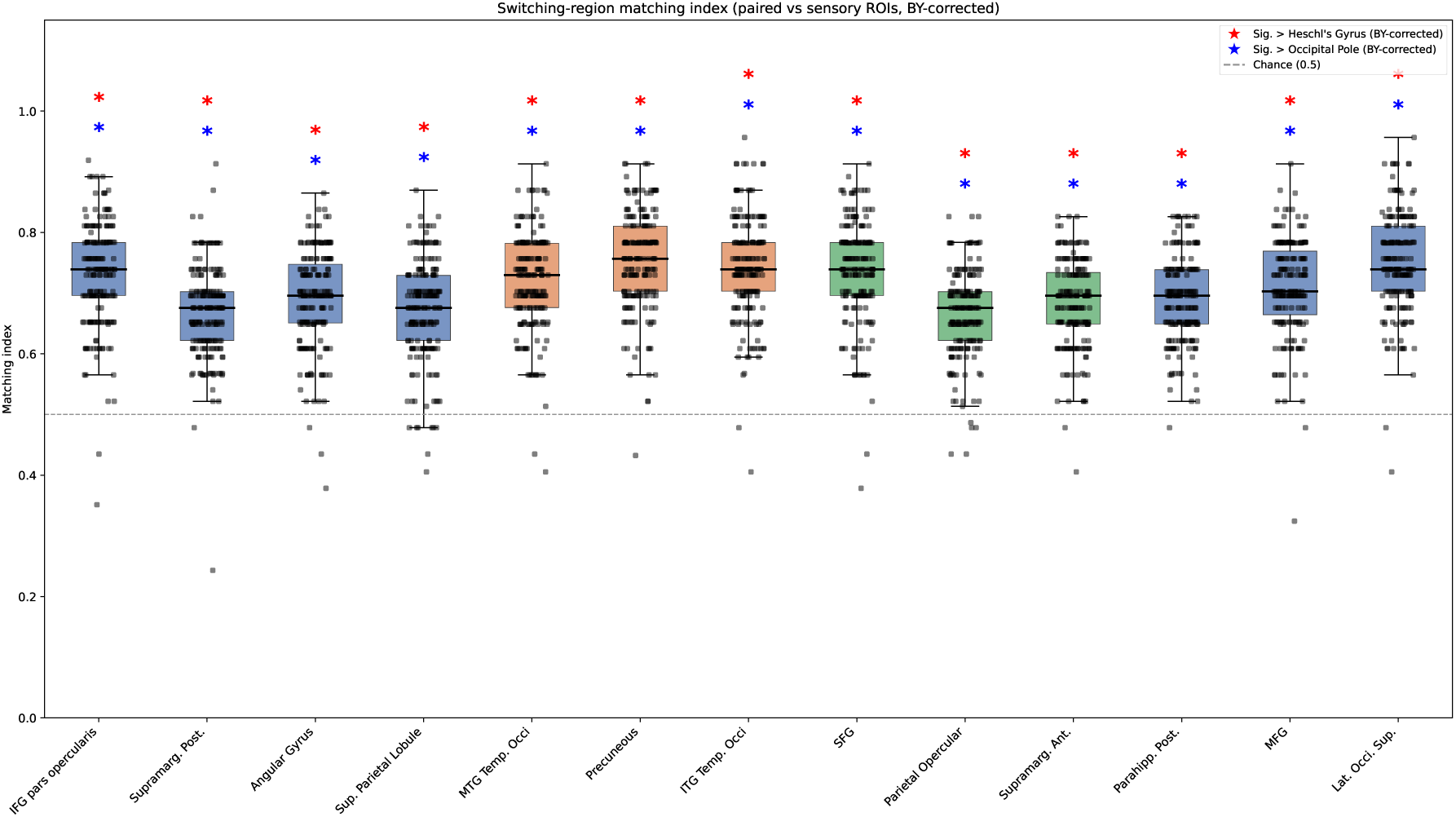
Matching index distribution for every participant recruited on the behavioral experiment across the switching regions. A paired two-sample permutation test was done for each switching region to establish if the matching index is higher for that region over each of the two sensory regions (Occipital Pole and Heschl’s Gyrus (H1 and H2)) and multiple testing correction was done using Benjamini-Yekutieli procedure. The matching indices were found to be significantly higher for all ROIs than the sensory regions.

**Extended Figure E5:**
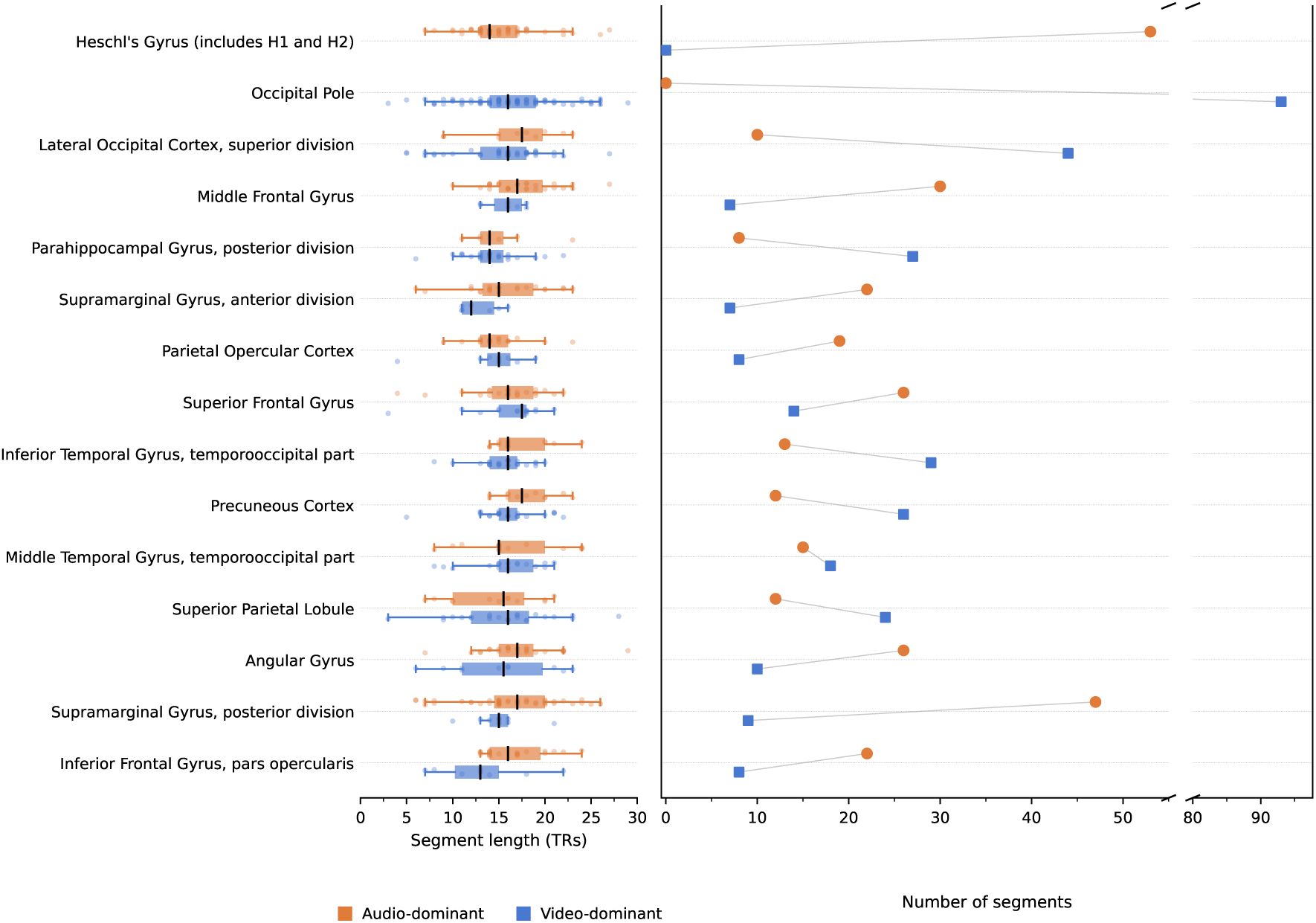
Left, distribution of audio- and video-dominant segment lengths for each switching region, with Heschl’s gyrus and the occipital pole shown as auditory and visual baselines. Right, number of audio- and video-dominant segments per region; lines connect the two counts for each region for visual clarity. Dominant segments in switching regions are of intermediate length, mostly spanning 10–18 TRs (*∼*15–27 s). Each switching region therefore favours each modality across multiple scenes of intermediate duration, rather than in a few prolonged stretches or brief transients.

**Extended Figure E6:**
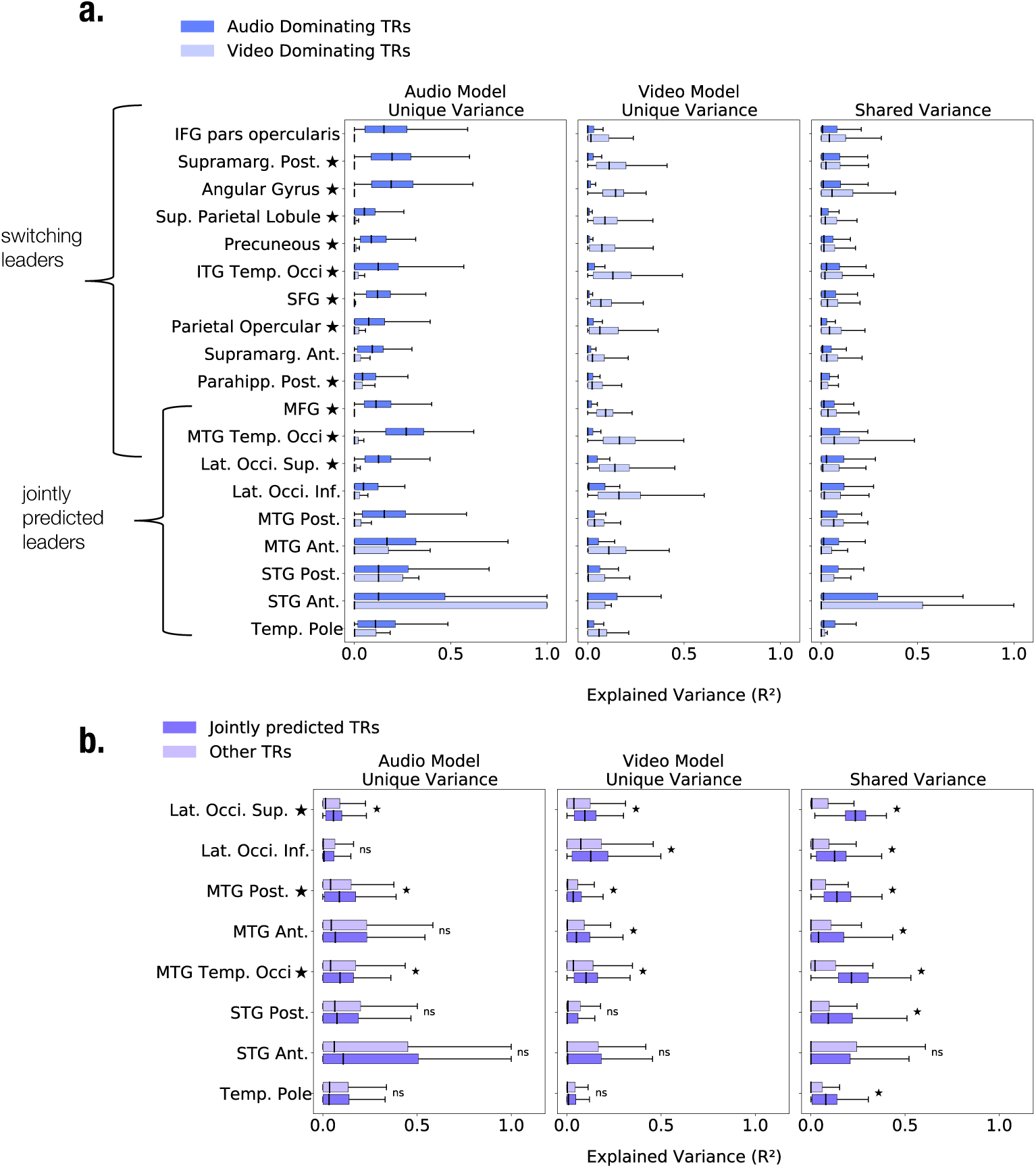
(a) Variance-decomposition results for switching and jointly-predicted leaders during audio-dominating and video-dominating segments (DIP). Box plots show distribution across TRs of unique variance(R2) explained by audio and video features, and of shared variance. Boxes span the interquartile range (IQR) with the median marked, whiskers extend to 1.5*×*IQR. Asterisks mark regions where the dominant-modality-unique variance significantly exceeds other components (Paired t-test, corrected at p ¡0.05 across all tests). Regions up to and including lateral occipital superior are switching leaders, the remaining ones including MTG temp. occi. and lateral occipital superior are jointly predicted. Most switching regions show significant modality-unique dominance, indicating reliance on the prevailing sensory modality, whereas jointly-predicted regions maintain more balanced encoding even during unimodal-dominant segments. (b) Variance components for jointly-predicted leaders during jointly-predicted TRs (labeled both) versus the remaining TRs (labeled other). Asterisks next to region names indicate that shared variance significantly exceeds both unique components (Wilcoxon paired t-test, corrected at p ¡ 0.05 across all tests), and asterisks on horizontal bars denote that the corresponding variance component during jointly-predicted TRs is significantly greater than that during the rest. All divisions of MTG and Lat. Occi. retain significant modality-unique components, suggesting that even within periods of high audiovisual joint-prediction, these regions encode complementary modality-specific information.

**Extended Figure E7:**
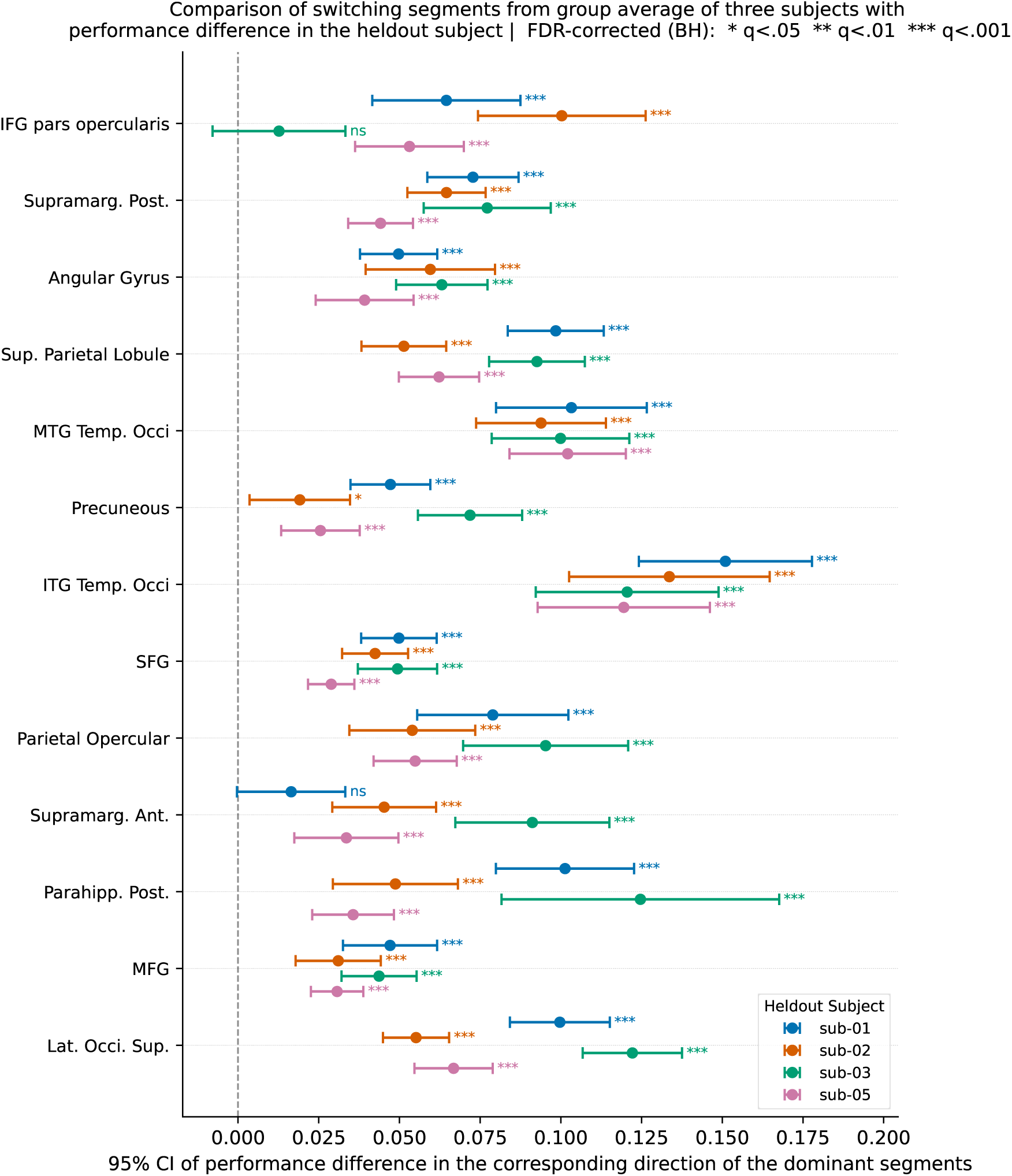
Directional overlap of predictive performance on one subject with switching segments from the group average of remaining subjects. For the regions along the y-axis, we compute the switching segments using averaged data from three subjects and plot the 95% CI of the performance difference of the remaining subject in the corresponding direction (audio - video performance for audio segments and video-audio performance for video segments) and use a one sample t-test to see if the performance difference is significantly above zero and correct for multiple testing. 50 out of 13 *×* 4 = 52 CIs are above zero, indicating substantial cross-subject overlap.

**Extended Figure E8:**
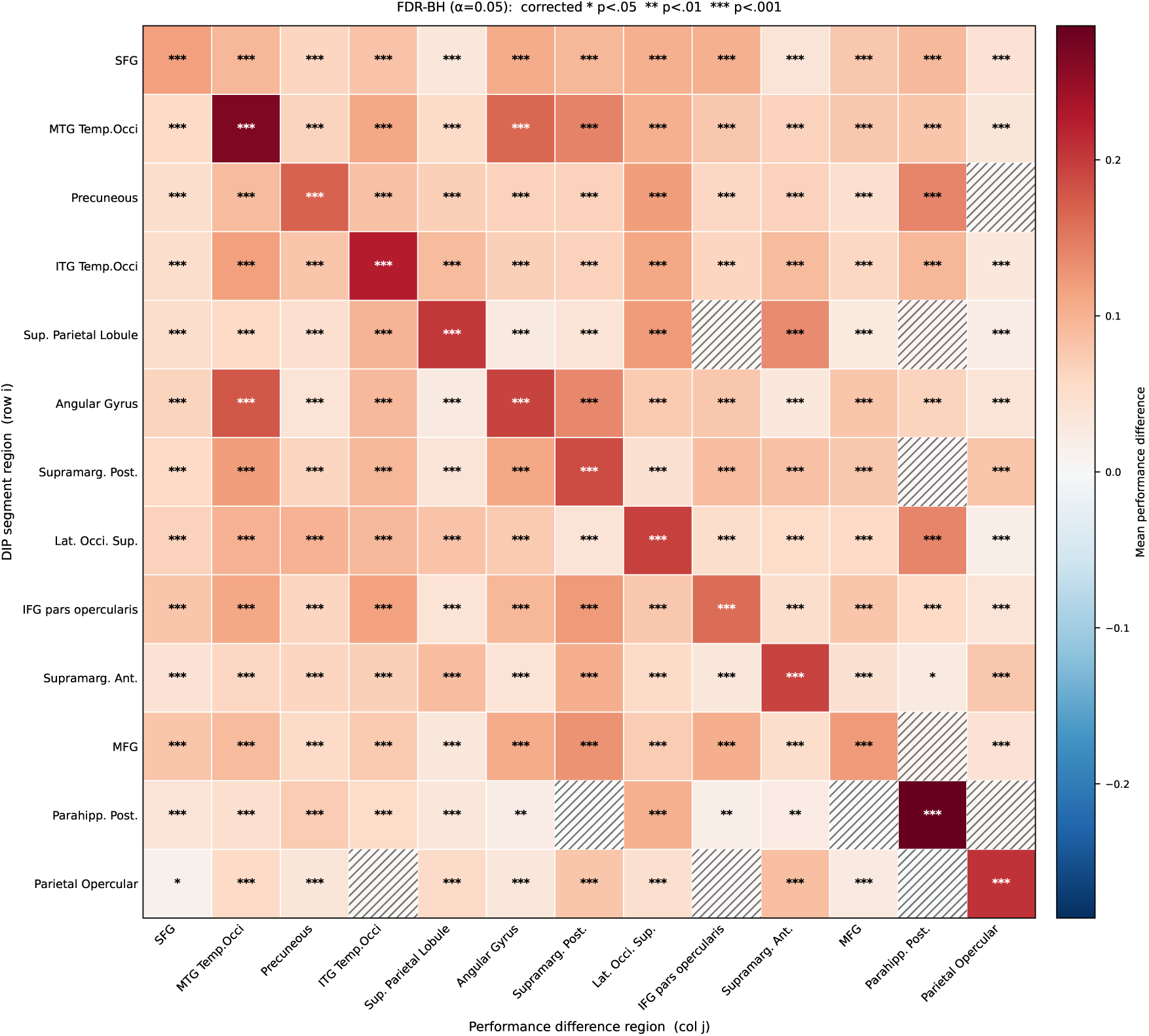
Directional overlap of predictive performance on a region with switching segments from every other switching region. Rows denote the regions used to define audio- and visual-dominant segments, whereas columns denote the regions whose performance difference was evaluated on those segments in the corresponding direction. So for audio- and video-dominant segments for a region along the y axis, the respective performance difference (audio-video and video-audio) for those segments was computed for every region along the x-axis. Each cell shows the mean performance difference, and significance of performance difference being above zero was determined using a one-sample t-test, with significance levels annotated on the plot after multiple testing correction. The significance test was done to show if the performance difference in the corresponding direction is significantly above zero, which would indicate strong directional overlap. Note that 145/156 non-diagonal cells of this matrix show significance, indicating strong directional overlap between ROIs. 10 out of the 11 cells that don’t show any significance are in the rows or columns of parahippocampal posterior or parietal opercular, indicating potentially more endogenous switching drive specific to these regions.

**Extended Figure E9:**
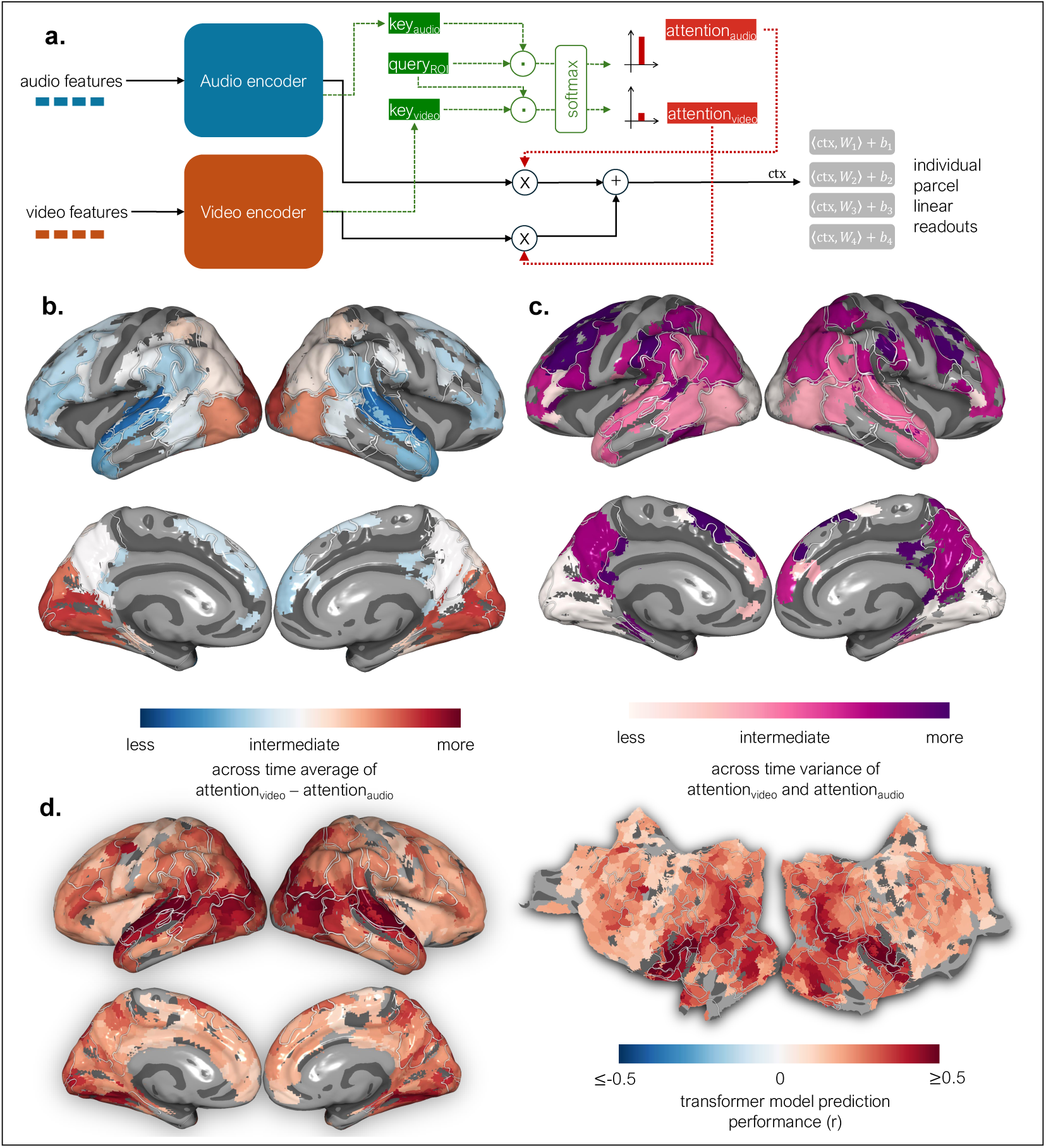
**(a)** Schematic of the ROI-specific audiovisual transformer used to predict parcelwise BOLD responses. A query matrix with ROI-specific learnable parameters determines the relative attention assigned to each modality as a function of the current embeddings for a specific region which makes the model learn how much weight to assign to audio versus video features independently for each ROI and timepoint. **(b)** Mean difference across time between attention weights for video and audio features (attention_video_ *−* attention_audio_) for every parcel. Positive values (red) indicate regions where the transformer consistently assigns a higher weight to video inputs, primarily around the occipital lobe. Negative values (blue) indicate higher weights for audio inputs, concentrated in superior and anterior temporal areas. Regions with intermediate mean differences can reflect two distinct cases: (1) the model alternates between assigning high weight to one modality and then the other across time, yielding an average near zero but reflecting temporal switching, or (2) the model maintains a balanced weighting of both modalities throughout, indicating sustained joint attention. **(c)** Temporal standard deviation of the attention-weight difference (attention_video_ *−* attention_audio_). High variance (magenta) indicates strong temporal fluctuations in relative attention, often co-localized with switching regions in frontal cortex and precuneus. Low standard deviation (light pink) indicates stable weighting of the two modalities across time, characteristic of unimodal regions focusing on one modality and jointly-attentive regions that maintain a consistent balance between auditory and visual information. **(d)** Pearson correlation performance of the model in held-out data.

