## Supplemental material referred to in the manuscript text for "Two cortical mechanisms for natural audiovisual processing"

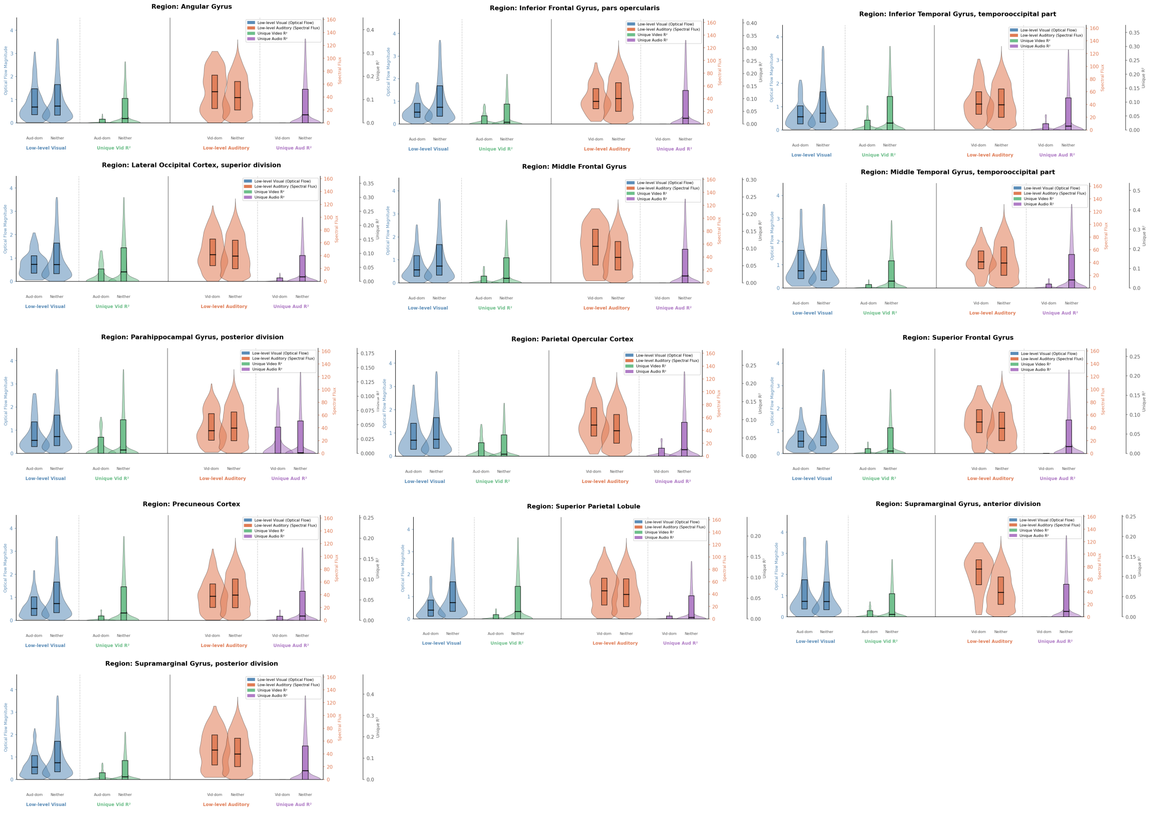


**Figure S1:** For each switching region, on the left panel, we show the distribution of low-level visual features and $R^2$ uniquely explained by video for audio-dominating TRs and neither TRs, and on the right panel, we show the distribution of low-level auditory features and $R^2$ uniquely explained by audio for video-dominating TRs and neither TRs. For most switching regions, the drop in explained unique variance by the non-dominant modality during the opposite modality dominant segments relative to usual TRs (TRs that are neither audio nor video dominating and are referred to as "neither" TRs in the plot) is substantially more than the drop in low-level features of the non-dominant modality, if that exists at all.

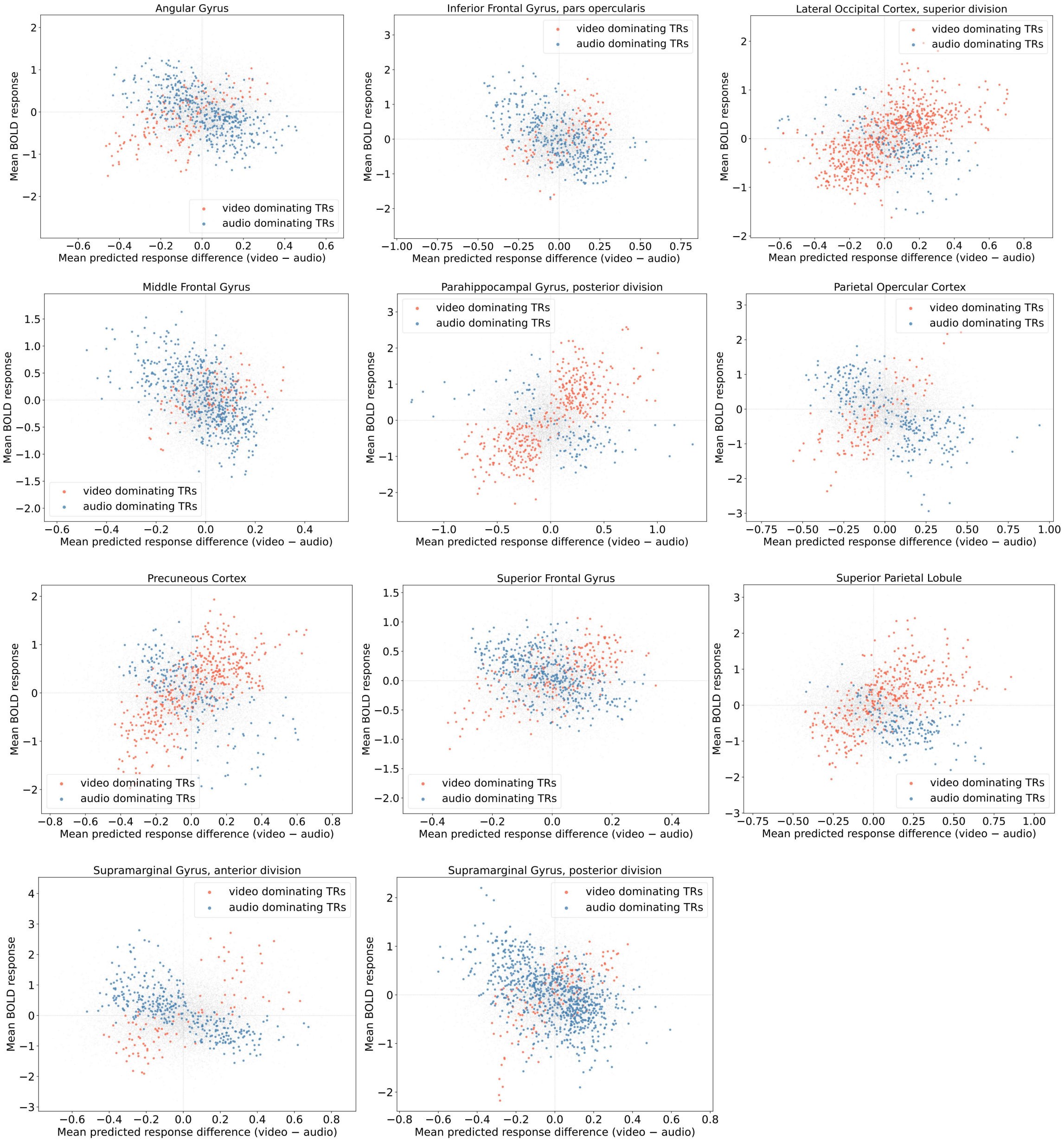


**Figure S2:** All switching regions with BOLD response averaged across parcels in a TR (along the y-axis) vs predicted response difference between audio and video embedding fitted model (video-audio, along the x-axis). The audio and video dominating TRs are respectively colored with blue and orange in the plot.

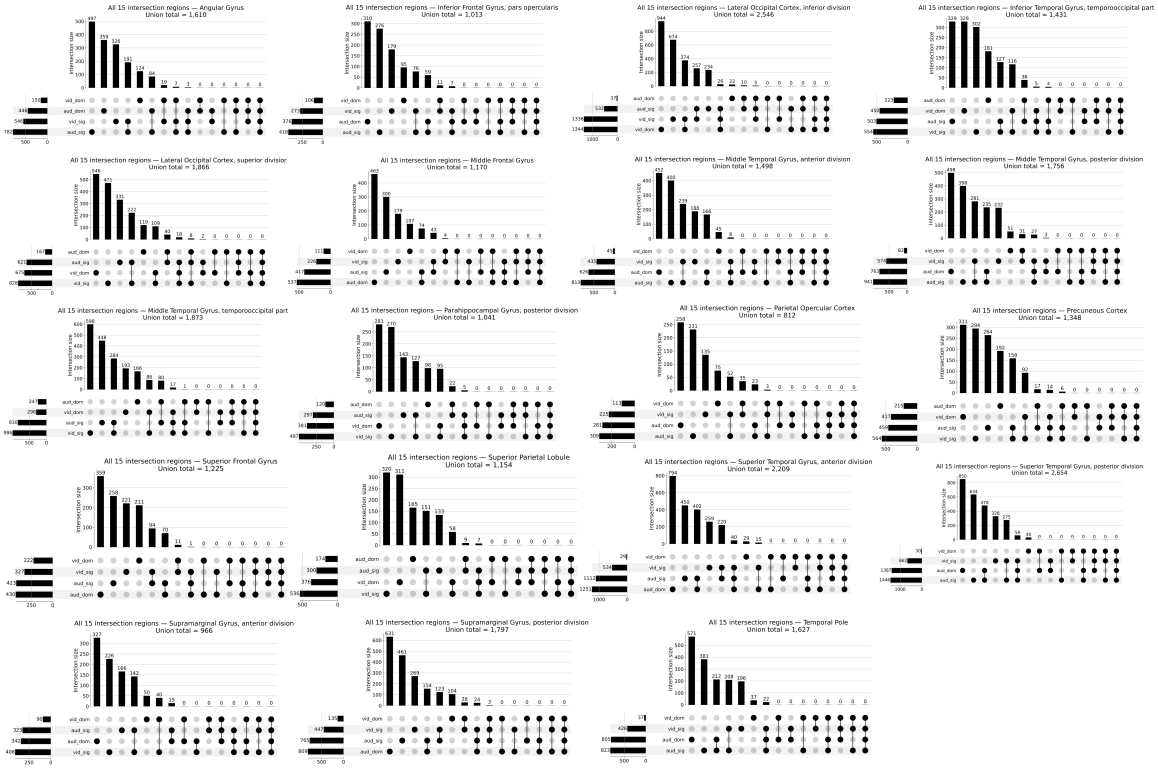


**Figure S3:** Following identification of audio-dominant and video-dominant segments via DIP, and highly predictable segments for the audio and video modalities via PIP (highly predictable as defined in Methods), we visualize the 15 non-empty subsets determined by a 4-set union for all switching and jointly predicted regions. For brevity, we refer to these highly predictable segments as "aud_sig" and "vid_sig" and the dominating segments as "aud_dom" and "vid_dom". The upset plots' bars are also ordered according to the size of the sets, so the order is not the same for every plot.

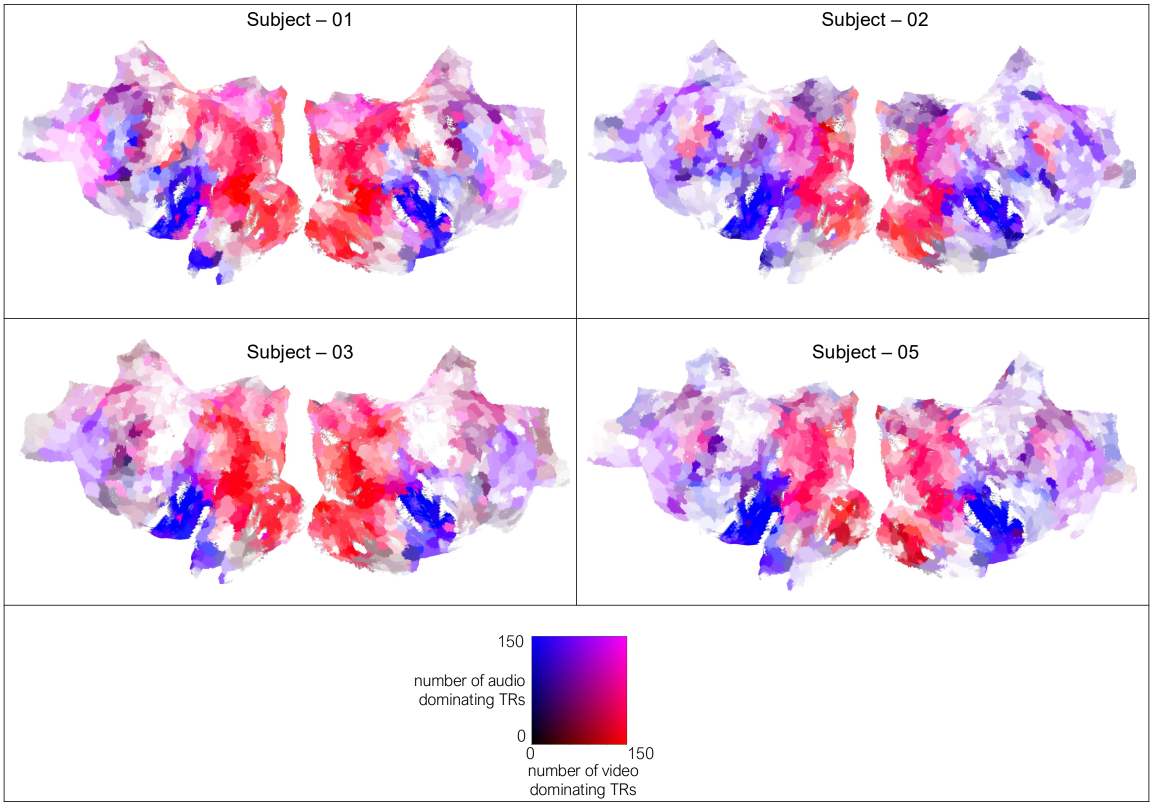


**Figure S4:** Spatial motif of switching regions in every subject when the trial average of a small subset of the stimuli with subjectwise repeats is considered. The color panel works similarly to that reported in the main text, Figure 3(a).

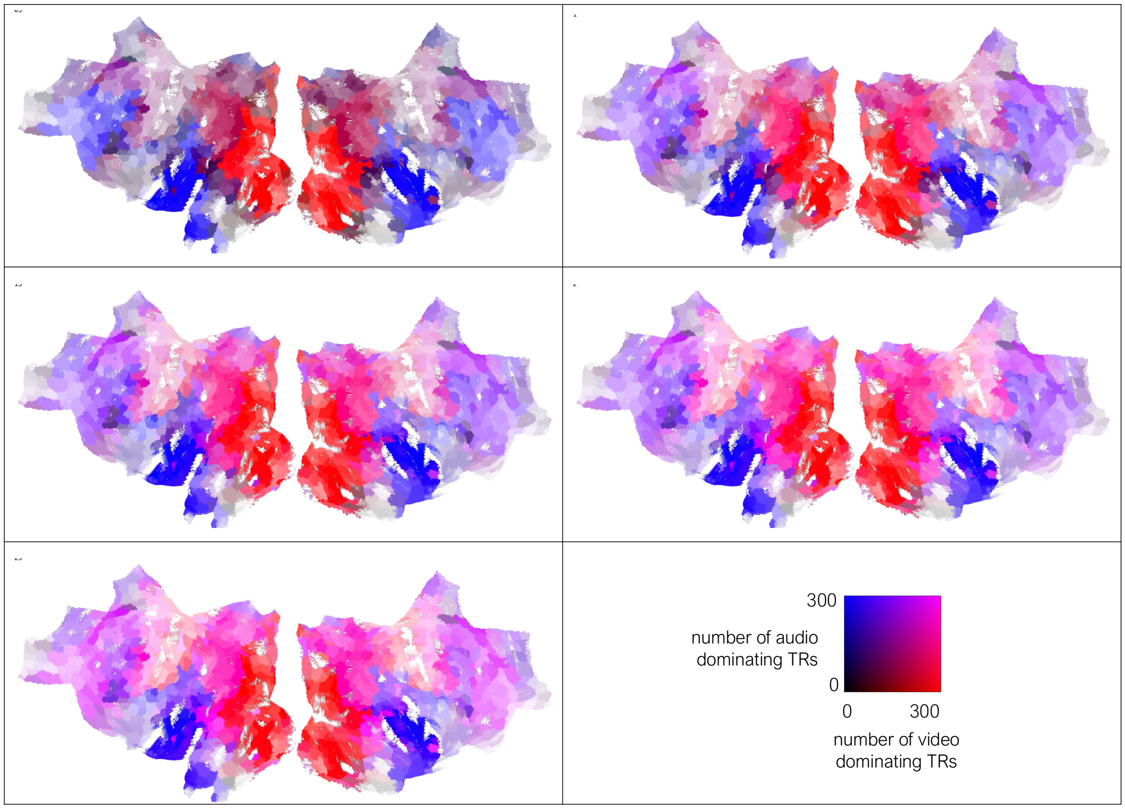


**Figure S5:** The left-to-right, starting from the top row, shows the spatial motif of switching regions for thresholds of 95%, 90%, 85%, 80%, and 75%. The brain flat maps were made using the same color scale and scheme as shown in Figure 3(a) of the manuscript. We think that the trend of switching in two bows becomes very evident from the 90% threshold and only becomes slightly clearer (clear as in visually convincing purple/magenta/pink color indicating a decent number of dominant TRs from both modalities) with a more lenient threshold. The frontal bow is pretty clear even for a 95% threshold.

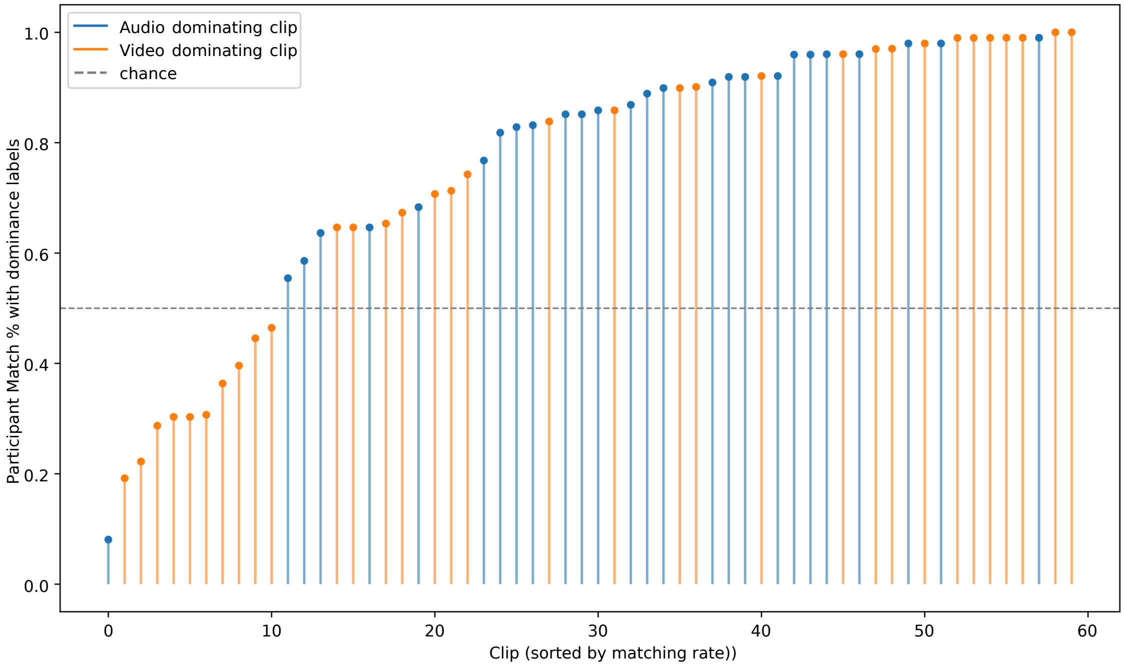


**Figure S6:** Matching rate between behavioral ratings from participants and neural dominance labels for every clip, ranked in increasing order and colored by dominance label from DIP.
